# Molecular basis for condensin enrichment at pericentromeres

**DOI:** 10.1101/2024.03.27.586992

**Authors:** Menglu Wang, Juan Zou, Christos Spanos, Juri Rappsilber, Adele L. Marston

**Affiliations:** The Wellcome Centre for Cell Biology, Institute of Cell Biology, University of Edinburgh, Edinburgh EH9 3BF, United Kingdom; Institute of Biotechnology, Technische Universität Berlin, Gustav-Meyer-Allee 25, 13355 Berlin, Germany

## Abstract

Faithful chromosome segregation requires packaging of the genome on both global and local scales. Condensin plays a crucial role at pericentromeres to resist spindle forces and ensure the bioriented attachment of kinetochores to microtubules in mitosis. Here we demonstrate that budding yeast condensin is recruited to pericentromeres through a direct interaction between its Ycg1 subunit and the pericentromeric adaptor protein, shugoshin (Sgo1). We identify a Short Linear Motif (SLiM), termed CR1, within the C-terminal region of Sgo1 which inserts into a conserved pocket on Ycg1. Disruption of this interface abolishes the Sgo1-condensin interaction, prevents condensin recruitment to pericentromeres and results in defective sister kinetochore biorientation in mitosis. Similar motifs to CR1 are found in known and potential condensin binding partners and the Ycg1 binding pocket is broadly conserved, including in the mammalian homolog CAP-G. Overall, we uncover the molecular mechanism that targets condensin to define a specialized chromosomal domain.

## Introduction

Accurate chromosome segregation protects against aneuploidy and requires precise packaging of the genome. Mitotic chromosome architecture is defined by an interplay between the Structural Maintenance of Chromosomes (SMC) complexes cohesin and condensin, which extrude DNA loops^1,2^. Cohesin has gained the additional property of tethering the sister chromatids together during mitosis to provide the cohesion to resist microtubule-dependent pulling forces^1^. In all SMC complexes, the molecular motor is a dimer of SMC proteins whose ATPase heads are bridged by kleisin proteins^1^. Association of the SMC core complex with a number of accessory proteins modulates their activity and targeting to confer specific functions in looping and/or cohesion. In the case of cohesin and condensin, these accessory proteins are part of a family of hook-shaped HEAT-repeat proteins, termed HAWKs^3^. Cohesin HAWKs are Scc3/SA2 and the inter-changeable Scc2/NIPBL and Pds5 proteins^1^. Yeast have a single condensin with HAWKs Ycg1 and Ycs4, while vertebrates have condensin I and II, with distinct HAWKs (CAP-G/CAP-D2 or CAP-G2/CAP-D3). In vertebrates, cohesin initiates chromosome separation in interphase^4^, however, most cohesin is removed during prophase, leaving only the pericentromeric pool holding sister chromatids together^5^. Compaction during prophase is driven by condensin II which generates long loops that are further divided into shorted nested loops after nuclear envelope breakdown by condensin I^6^. In yeast, cohesin drives the majority of chromosome compaction in mitosis, while condensin has specific roles at the rDNA and pericentromeres^7,8^.

At metaphase, mitotic chromosomes attach to microtubules from opposite poles via their sister kinetochores. Kinetochores reside on centromeres, which are embedded within larger chromosomal domains called pericentromeres, whose architecture is defined by cohesin and condensin^9–14^. Pericentromeres perform critical structural and regulatory roles in chromosome segregation including orienting kinetochore-microtubule attachments, resisting spindle forces and engaging surveillance mechanisms to sense improper attachments^15,16^. Cohesin enrichment at pericentromeres prevents the premature separation of sister centromeres and allows tension establishment^17–20^. Condensin organises vertebrate centromeres into two domains^10^ and is important in regulating centromere stiffness^9^. Consistently, condensin protects against merotely (where a single sister kinetochore is captured by microtubules from opposite poles)^11^.

Initial insights into how SMC proteins are specifically targeted to chromosomal loci to define their architecture came from budding yeast pericentromeres. Their “point” centromeres are defined by a short ∼125bp core sequence upon which the kinetochore assembles^15^. A phosphorylation-dependent interaction between the Ctf19 kinetochore protein and a conserved patch on the Scc4 component of the Scc2-Scc4 cohesin loader targets cohesin to the core centromere^21,22^. Centromere-loaded cohesin extrudes a loop on either side to shape the ∼10kb pericentromere, and convergent genes flanking centromeres set the position of the pericentromere borders by acting as barriers to loop extrusion and sites of sister chromatid cohesion^20^. This loop-based structure at pericentromeres provides a platform for surveillance mechanisms that direct and sense the establishment of biorientation^23^. The pericentromeric adaptor protein, Shugoshin, Sgo1, acts downstream of cohesin and plays a key role in this process. Sgo1 facilitates sister kinetochore biorientation by promoting the pericentromeric recruitment of a number of effectors including PP2A, Aurora B and condensin^24–26^. Condensin biases sister kinetochores towards biorientation in early mitosis and promotes the topoisomerase II-dependent removal of residual centromeric catenanes^24,27^. Following the establishment of biorientation, sister kinetochore tension leads to dissociation of Sgo1, condensin and loop-extruding cohesin from pericentromeres, while cohesive cohesin is retained at pericentromere borders to resist spindle forces^18,20,28–30^.

Specific targeting of SMC complexes through their HAWK subunits may be a general principle underlying the architecture of specialized chromosomal domains. The conserved patch on the cohesin loader Scc4 that binds Ctf19 also engages with the RSC chromatin remodeler complex and may contribute to cohesin loading genome wide^31^. Similarly, the equivalent patch on the Scc4 ortholog, MAU2, mediates cohesin recruitment to chromosomes in *Xenopus* extracts^21^ and a small deletion of this region in human MAU2 causes the developmental disorder Cornelia de Lange syndrome^32^. Although the molecular basis of condensin targeting is unknown, a number of candidate receptors have been described, in addition to yeast Sgo1^24,25,33^. In fission yeast and *Candida albicans*, monopolin proteins (Lrs4-Csm1/Mde4-Pcs1) promote the centromeric enrichment of condensin and to prevent merotely^34–36^. Similarly, monopolin (Lrs4-Csm1) is important for condensin localization and function within the rDNA and mating locus in budding yeast^37,38^. In *Xenopus* egg extracts, the general transcription factor TFIIH promotes the enrichment of condensin on chromatin, though this may be indirect^39^. In vertebrates, a short sequence in the tail of the chromokinesin Kif4A is implicated in condensin regulation and recruitment to chromatin^40–44^.

Here we reveal that condensin is recruited to budding yeast pericentromeres through a direct interaction between a Short Linear Motif (SLiM) on Sgo1, termed Conserved Region 1 (CR1), and a conserved patch on the HAWK subunit Ycg1. Mutation of specific residues in Sgo1 or Ycg1 that disrupt this interface demonstrates the importance of pericentromeric condensin for sister kinetochore biorientation in mitosis. Importantly, we find that the Ycg1 patch that recruits condensin to pericentromeres is conserved throughout evolution, including in the human ortholog, CAP-G and identify CR1 motifs in other candidate condensin interactors. Our findings suggest a general mechanism by which condensin, and potentially other SMCs, interact with ligands involved in its targeting and regulation.

## Results

### Condensin interacts directly with Sgo1 through its Ycg1 subunit

We previously found that condensin subunits co-immunopurify with Sgo1 in metaphase-arrested yeast extracts^24,45^. To understand whether Sgo1 binds to condensin directly, we purified full length Sgo1 and tested its ability to interact with the pentameric condensin holocomplex (Figure 1A). Immobilized V5-Sgo1 on beads specifically pulled down the condensin holocomplex (Figure 1B and C). We also found that the presence of Sgo1 caused condensin to elute at a lower volume in analytical SEC, providing further evidence for a direct interaction (Figure S1A). To determine which subunit of condensin Sgo1 interacts with, we performed Crosslinking Mass Spectrometry (CLMS). We first validated our CLMS data by mapping of inter-condensin subunit crosslinks onto a previously published cryo-EM structure of tetrameric condensin (lacking Ycg1)^46^ (Figure S1B: Table S1), which confirmed the detection of *bona-fide* interactions. Restricting our analysis to crosslinks involving Sgo1 revealed that the majority of crosslinks mapped to Ycg1, with a small number mapping to Smc4 in a region known to be close to Ycg1^46^ (Figure 1D). The largest number of crosslinks were in the Sgo1 C terminal region, with a smaller number mapping to an N terminal region (Figure 1D). This suggested that a major Ycg1 binding site resides in the C terminal region of Sgo1, while a minor binding site may exist in the N terminal region of Sgo1. To address this more directly, we tested whether recombinant Sgo1 N- or C-terminal fragments could associate with condensin from yeast extracts. Extracts were generated from cells lacking endogenous Sgo1 to preclude multimerization, and condensin binding to Sgo1 fragments immobilized on beads was detected by immunoblotting against Brn1-6HA (Figure S1C). Consistent with the CLMS, this indicated that the C-terminal region of Sgo1 (residues 350-590) harbors the predominant condensin binding site but that there is an additional minor binding site in the N-terminal region (residues 1-130). A previous report using peptide arrays suggested that Sgo1 harbors a condensin-binding region within its Serine-Rich Motif (SRM, residues 137-163)^33^, however, in our hands a recombinant fragment (130-190) containing the Sgo1 SRM did not pull down condensin (Figure S1C). Since Sgo1(350-590) showed the most robust condensin association, we next asked if this region is sufficient for binding to the pure pentameric condensin complex (Figure 1E). Reciprocal immunoprecipitation of condensin tagged with Brn1-6HA and V5-Sgo1 revealed that V5-Sgo1(350-590) itself bound non-specifically to anti-HA beads, preventing assessment of its ability to bind to immobilized condensin (Figure 1E). However, immobilized V5-Sgo1(350-590) specifically pulled down pentameric condensin, indicating that the C terminal region of Sgo1 (residues 350-590) is sufficient for condensin binding (Figure 1E). We conclude that Sgo1(350-590) binds condensin directly through the Ycg1 subunit.

**Figure 1.**
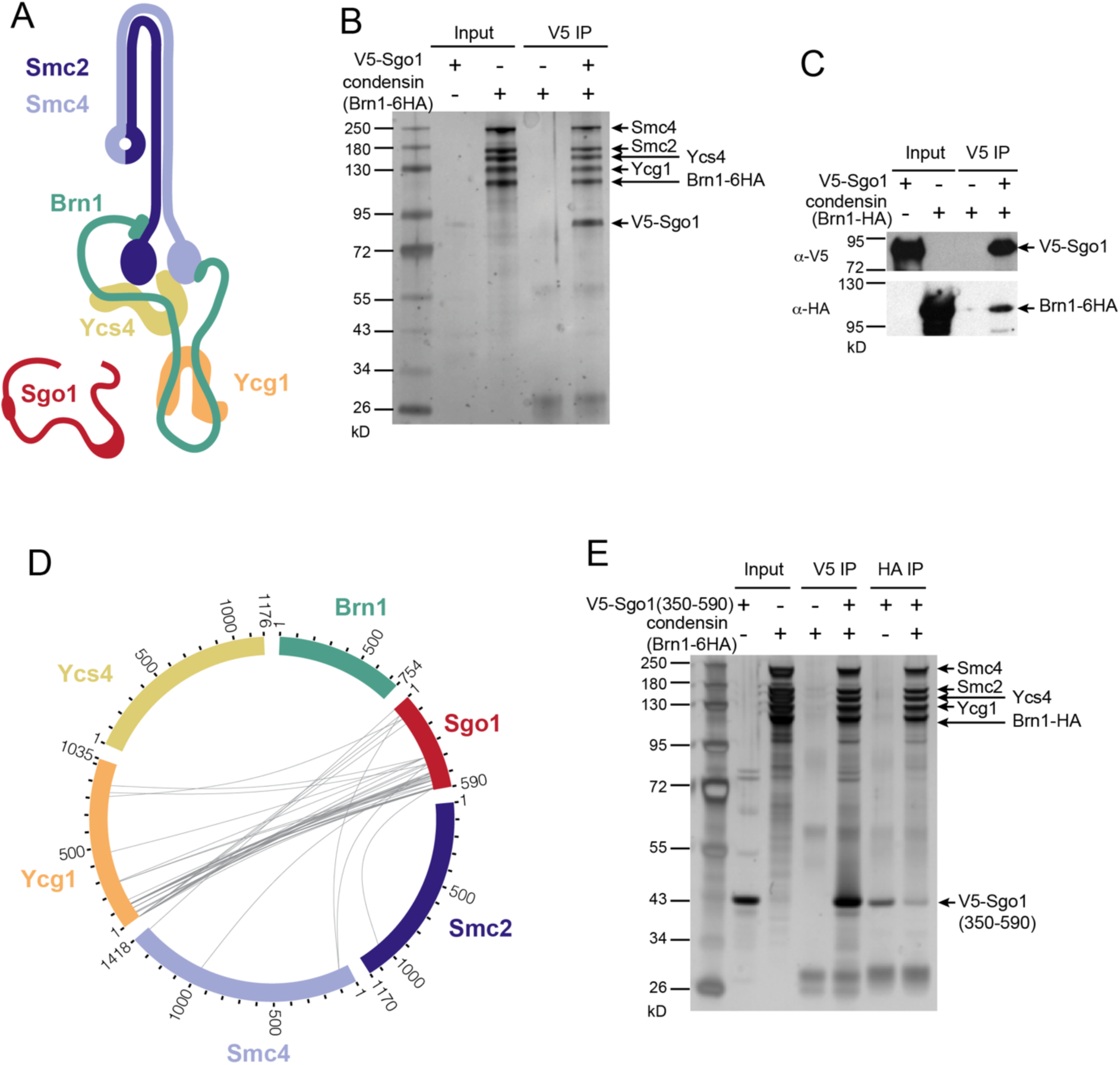
Sgo1 and condensin interact directly. (A) Schematic representation of Sgo1 and condensin complex. (B, C) Recombinant *Sc* Sgo1 co-immunoprecipitated with *Sc* condensin complex using anti-V5-coupled beads. Immunoprecipitates were analyzed by silver-stained SDS-PAGE (B) and immunoblotted with the indicated antibodies (C). (D) Crosslinking mass spectrometry of full length Sgo1 and condensin holocomplex using crosslinker BS3. Intra-subunit condensin interactions and self-links were hidden and only crosslinks with score>10.5 are shown. (E) Co-immunoprecipitation assay using recombinant C-terminus of Sgo1 (350-590) and condensin complex, eluates were visualized by silver-stained SDS-PAGE gel. See also Figure S1. Experiments in Figure 1B,C and E performed once as shown.

### CR1 - a SLiM in the C terminal region of Sgo1 mediates Ycg1 binding

To identify the determinants of the Sgo1-Ycg1 interaction we purified Ycg1(6-932, Δ499-555)-Brn1(384-529), as described previously^47^, and generated a complex with Sgo1(350-590) (Figure 2A; Figure S2A). We performed CLMS and again validated our results by mapping crosslinks onto the reported Ycg1(6-932, Δ499-555)-Brn1(384-529) crystal structure (Figure S2B; PBD:5OQQ)^47^ (Figure S2B; Table S1). Crosslinks between Ycg1(6-932, Δ499-555)-Brn1(384-529) and Sgo1(350-590) indicated a major interface between the C terminal region of Sgo1 (around or after residue 500) and the N terminal region of Ycg1 (Figure 2B).

**Figure 2.**
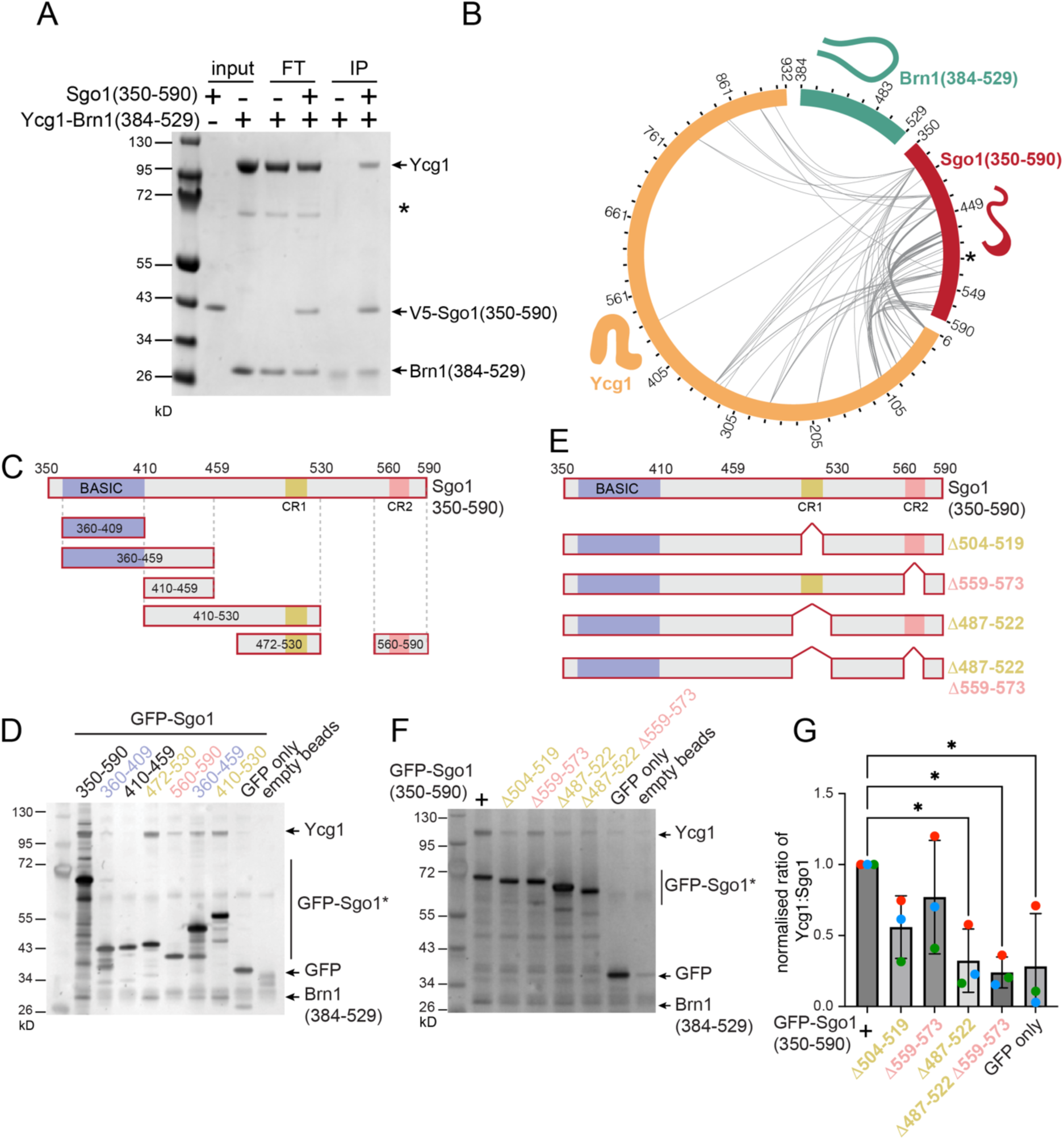
A conserved region within the C-terminus of Sgo1 is required for Ycg1 binding. (A) V5 tagged recombinant C-terminal region of Sgo1 (350-590) co-immunoprecipitated with Ycg1 (6-932, Δ499-555)-Brn1 (384-529). Eluates were analyzed by silver-stained SDS-PAGE. Asterisk indicates impurity, FT is flow through. (B) BS3 crosslinking mass spectrometry of Sgo1 (350-590) and Ycg1 (6-932, Δ499-555)- Brn1 (384-529). The interactions with Brn1 and self-links are hidden. Crosslinks with score>10.5 are chosen. Asterisk highlights Sgo1 residues (L508, F509) predicted to bind Ycg1. (C) Scheme showing the recombinant Sgo1 C-terminal variants generated in this work. The basic motif involved in Sgo1 chromosome localisation is highlighted in blue. Two conserved regions found by alignment of yeast sequences (Figure S3) are indicated in yellow and pink. (D) Recombinant GFP-tagged C-Sgo1 variants (coloured according to Figure 2C) co-immunoprecipitated with Ycg1 (6-932, Δ499-555)-Brn1 (384-529) using anti-GFP-coupled beads. Immunoprecipitates were analyzed by silver-stained SDS-PAGE. Performed once as shown. (E) Schematic diagram of the Sgo1 recombinant C-terminal deletion mutants used in the co-immunoprecipitation assay. (F, G) Recombinant GFP-tagged C-Sgo1 deletion mutants (coloured according to Figure 2E) co-immunoprecipitated with Ycg1 (6-932, Δ499-555)-Brn1 (384-529) using anti-GFP-coupled beads were analyzed by silver-stained SDS-PAGE (F). The ratios of Ycg1/Sgo1 gel bands intensity were calculated and the mean level from three experimental repeats with error bars representing standard deviation are shown in after normalization to wild type (G). *p<0.0332. One-way ordinary ANOVA with Dunnetts correction, only comparisons significantly different from wild type are indicated. See also Figure S2.

To further identify the potential binding site on Sgo1 we sought to identify conserved motifs that could indicate regions with critical functions. Despite significant functional conservation, shugoshins show a high degree of diversity^48^, therefore we restricted our analysis to related budding yeast species. These alignments of the C terminal region of Sgo proteins revealed that, in addition to the previously reported basic SGO motif that may associate with histones^49^, there are two further short <u>C</u>onserved <u>R</u>egions (CR1 and CR2) (Figure S3). Among a panel of recombinant Sgo1 fragments, we found that those containing the CR1 motif consistently pulled down Ycg1(6-932, Δ499-555)-Brn1(384-529) (Figure 2C and D). Furthermore, specific deletion of CR1, but not CR2, greatly reduced the ability of Sgo1(350-590) to pull down Ycg1(6-932, Δ499-555)-Brn1(384-529) (Figure 2E, F and G). Together, these experiments indicate that a major Ycg1 binding site resides with the CR1 motif of Sgo1.

The Sgo1 CR1 motif is characterised by an arrangement of hydrophobic residues (L508, F509, I513, V514 and L517 in *S. cerevisiae*) flanked by positively-charged lysine/arginine residues (Figure 3A). We mutated the Sgo1 CR1 hydrophobic residues to glutamic acid in various combinations and found that mutation of L508 and F509 had the strongest effect on Ycg1(6-932, Δ499-555)-Brn1(384-529) binding *in vitro* (Figure 3B and C). To address the importance of the CR1 motif *in vivo*, we adapted a tethering assay we developed previously^24,29^. We fused full length Sgo1, or its mutant derivatives, to TetR-GFP in cells carrying *tetO* arrays inserted at a chromosomal arm site (Figure 3D). To assay the ability of tethered TetR-GFP-Sgo1 to recruit condensin (Brn1-6HA) to this ectopic site we used chromatin immunoprecipitation, followed by qPCR (ChIP-qPCR). As expected, TetR-GFP-Sgo1 specifically recruited Brn1-6HA to the *tetO* arrays, however, mutations within Sgo1-CR1 reduced Brn1 recruitment (Figure 3E). Consistent with our *in vitro* analysis, the Sgo1-L508E F509E mutations had the strongest effect in abolishing Brn1 recruitment to this ectopic site (Figure 3E). We also found that deletion of the serine-rich motif (SRM) (Sgo1-Δ137-163) did not prevent Brn1 recruitment *in vivo* (Figure 3E), consistent with its dispensability for binding in our *in vitro* assay (Figure S1C), while contradicting expectations from a previous study^33^. Instead, our data indicate that L508 and F509 glutamic acid substitutions in the context of the full-length protein are sufficient to abrogate condensin association *in vivo.* Therefore, the Sgo1-Ycg1 interaction depends on the conserved adjacent hydrophobic residues L508 and F509.

**Figure 3.**
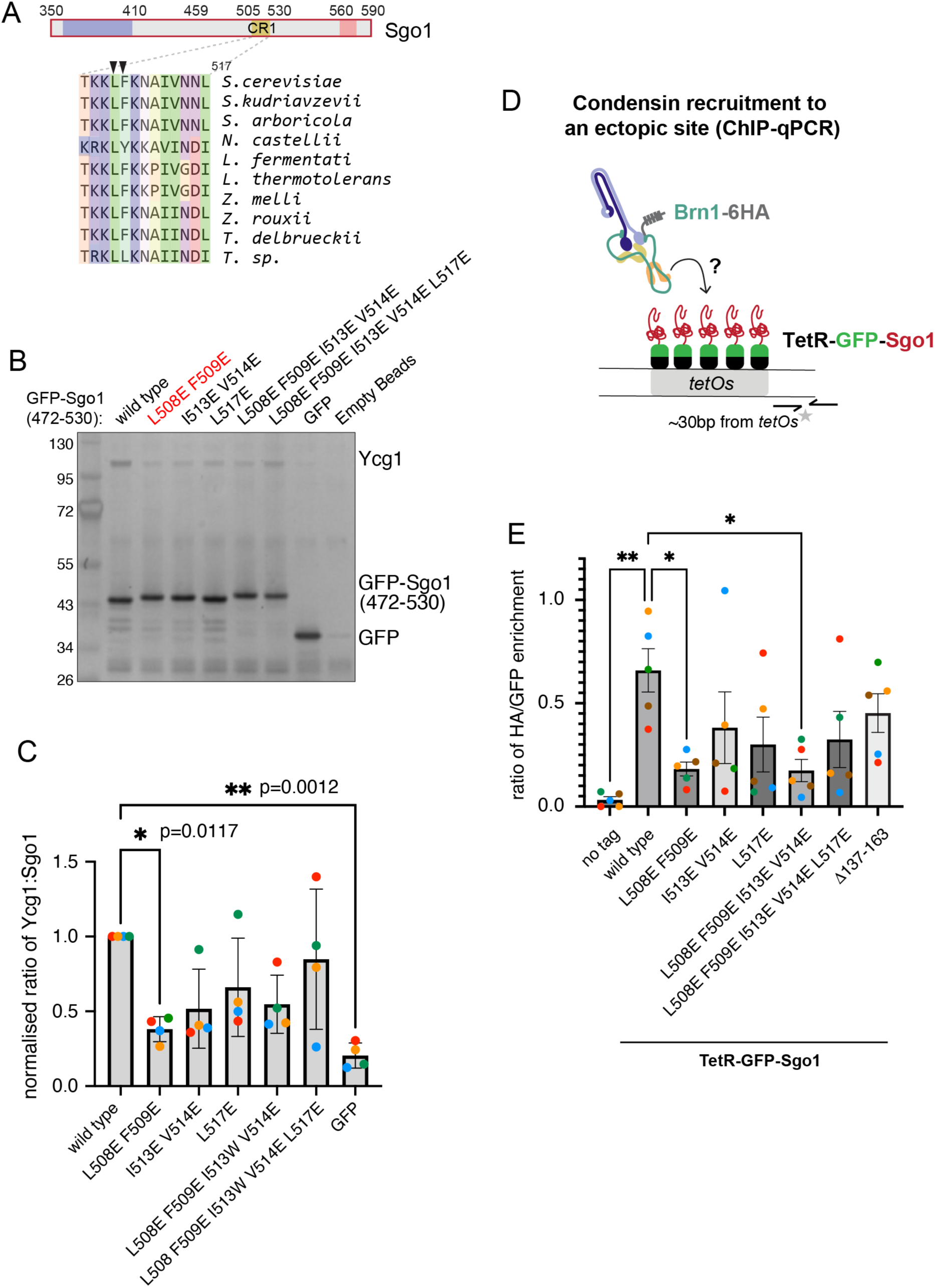
Mutation of Sgo1 Conserved Region 1 (CR1) disrupts condensin binding. (A) Alignment of sequences from related yeast species showing the short conserved region 1 on Sgo1. (B, C) Recombinant Sgo1 (472-530) point mutants co-immunoprecipitated with Ycg1 (6-932, Δ499-555)-Brn1 (384-529), the elutes were analyzed by silver stain (B) and the mean of Ycg1/Sgo1 gel bands intensity ratio from four experimental repeats are shown after normalization to wild type (C). Error bars represent standard deviation. *p<0.0332; **p<0.0021, one-way ordinary ANOVA with Dunnetts correction, only comparisons significantly different from wild type are indicated. (D) Schematic diagram showing tethered Sgo1 at an ectopic site recruits condensin, grey asterisk indicates the position of primers used for qPCR analysis, ∼30 bp distant from the tethering site. (E) ChIP-qPCR measuring the levels of Brn1-6HA recruited to the tethered wild type and mutant Sgo1 variants. Strains were arrested in mitosis by treatment with nocodazole and benomyl for 2 hours. The ratios of Brn1 and Sgo1 enrichment (anti-HA/anti-GFP ChIP-qPCR values were determined and the mean of four experimental repeats is shown with error bars representing standard error. *p<0.0332; **p<0.0021, one-way ordinary ANOVA with Dunnetts correction, only comparisons significantly different from wild type are indicated. See also Figure S3.

### Broad conservation of the predicted binding site in Ycg1/Cnd3/CAP-G condensin proteins

Next, we sought to identify the determinants on Ycg1 that mediate Sgo1 binding. We ran alphafold multimer predictions^50–52^ of Ycg1(6-932, Δ499-555)-Brn1(384-529) and Sgo1(487-522). Out of 5 highly similar models (Figure S4A and B). we considered Rank 5 to be the best predictor of the Sgo1-Ycg1 interface, since it reported the highest pLDDT scores for the two Sgo1 residues L508, F509 we identified to be important experimentally (Figure 3C and E). pLDDT is a reliable predictor of the Cα local-distance difference test^53^, providing confidence in the predicted position of the these residues (Figure S4C). All five models were consistent with our CLMS of full length Sgo1 with the pentameric condensin since distances between crosslinked residues on the models were all within the 25 Å reach of the BS3 crosslinker used, but shortest for Rank 5 (Figure 4A and B; Figure S4D and E; Table S2), indicating the best structural fit to our experimental data. The positions of residues flanking L508 F509 were also predicted with relatively high confidence, while the positions of more distant residues were not (Figure 4C). In all models, Sgo1 L508 and F509 fit into a hydrophobic binding pocket between HEAT repeats 3 and 4 on the concave surface under the nose of Ycg1 (Figure 4D). In the model, Sgo1 binds to a surface on Ycg1 that is opposite the Ycg1-DNA interface. Modelling Sgo1 onto the structure of Ycg1-Brn1 complexed with DNA further suggested that Ycg1-Brn1 could bind Sgo1 and DNA concurrently (Figure S4F). In the predicted structure, Sgo1 L508 inserts into a binding pocket comprised of Ycg1 residues M145, I148, I151, F156 and F185 (Figure 4D). Although Sgo1 CR1 is conserved only among yeasts closely related to *S. cerevisiae*, its predicted binding site on Ycg1 is highly conserved not only among yeasts (Figure S4G) but also among Ycg1/CAP-G proteins across phyla, including in mammals (Figure 4E), indicating it may provide a common binding surface for multiple ligands. We conclude that Ycg1/CAP-G proteins harbour a conserved hydrophobic pocket which provides a binding site for Sgo1 and potentially other ligands.

**Figure 4.**
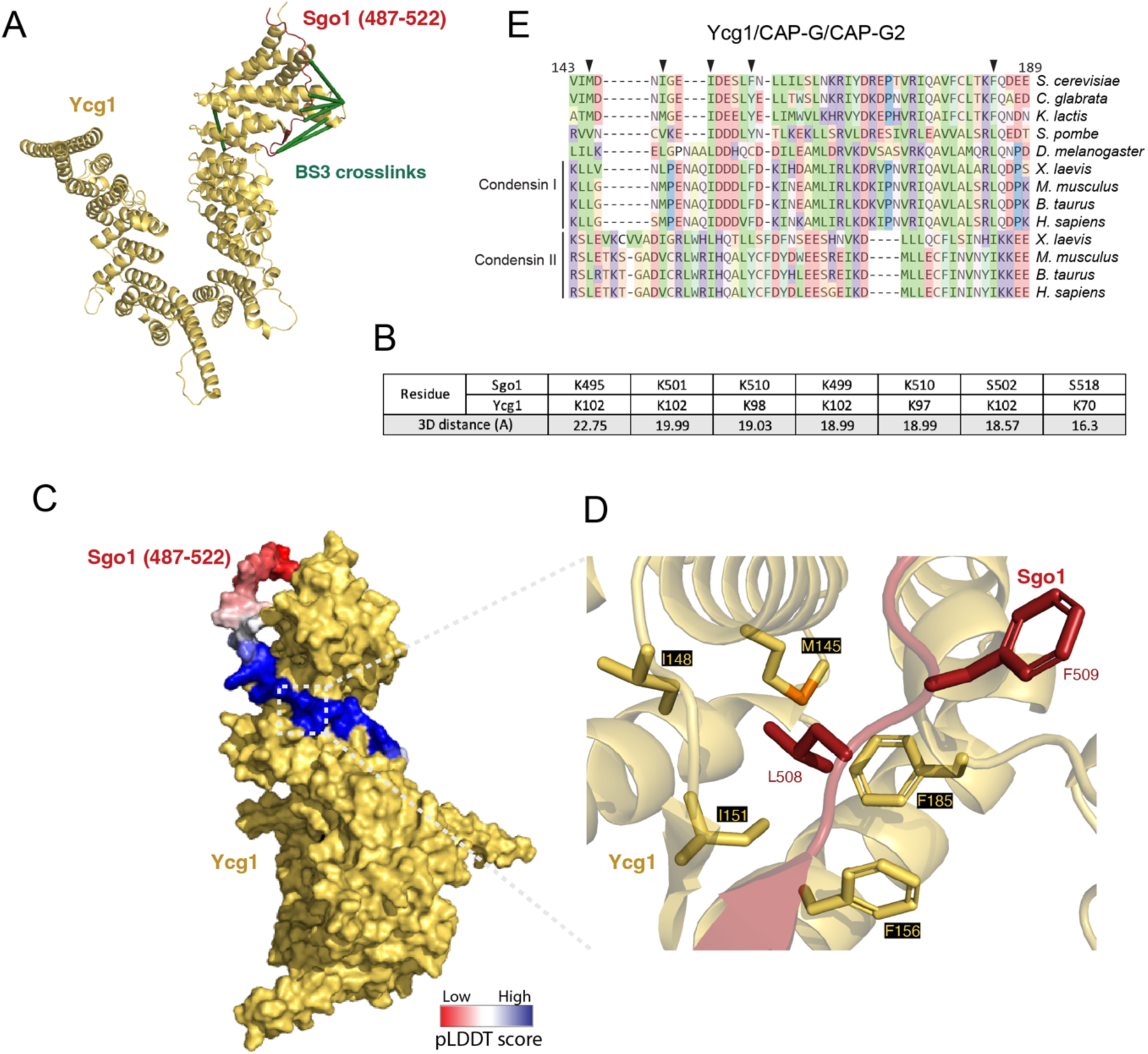
Identification of the Sgo1-Ycg1 interaction interface. (A) Mapping of crosslinks from CLMS of full length Sgo1 and condensin holo-complex data shown in Figure 1D onto AlphaFold2 model. (B) Table indicates distances between crosslinked residues on the AlphaFold2 model, with range of the BS3 crosslinker (25Å). (C,D) AlphaFold2 model of Sgo1(487-522) with Ycg1 (6-932, Δ499-555)-Brn1 (384-529), Sgo1 peptide (487-522) is coloured with pLDDT score. (D) with zoom in to show the detailed interaction surface between Sgo1 and Ycg1. (E) Conservation of the Ycg1/CAP-G binding pocket. Mutated residues are indicated with arrowheads. See also Figure S4 and S5.

### Mutation of the predicted Sgo1-Ycg1 interface disrupts their interaction *in vivo*

To determine whether the predicted Sgo1-Ycg1 binding site we identified through CLMS and alphafold is functionally important in mediating an interaction between the two proteins *in vivo*, we generated specific point mutations in the endogenous copies of *SGO1* and *YCG1* (Figure 5A). We mutated four residues M145, I148, I151 and F156 that form the hydrophobic Ycg1 pocket to alanine (*ycg1-4A*) and Sgo1 residues L508 and F509 to either alanine or aspartic acid (*sgo1-2A* and *sgo1-2E*, respectively). To understand the impact of these mutations on the Sgo1-Ycg1 interaction *in vivo*, we immunoprecipitated Sgo1 from metaphase-arrested cells and identified interacting proteins by mass spectrometry (Figure 5B-F; Figure S5). In Sgo1 immunoprecipitates from wild-type cells, all 5 condensin subunits were abundant, and we additionally identified components of the kinetochore, cohesin and PP2A, as expected (Figure 5B). However, comparison of Sgo1 immunoprecipitates from wild type with those from *sgo1-2A, sgo1-2E* or *ycg1-4A* cells revealed specific depletion of condensin subunits (Figure 5B-F), confirming the importance of the interface we identified (Figure 4D) in Sgo1-Ycg1 interaction. Importantly, similar amounts of Sgo1 were immunoprecipitated in all conditions (Figure S5A). Furthermore, other Sgo1 interactors, including PP2A and cohesin subunits, were not depleted in *sgo1-2A*, *sgo1-2E* or *ycg1-4A* cells compared to wild type (Figure S5B and C). Therefore, *sgo1-2A*, *sgo1-2E* and *ycg1-4A* mutations specifically abrogate the Sgo1-Ycg1 interaction.

**Figure 5.**
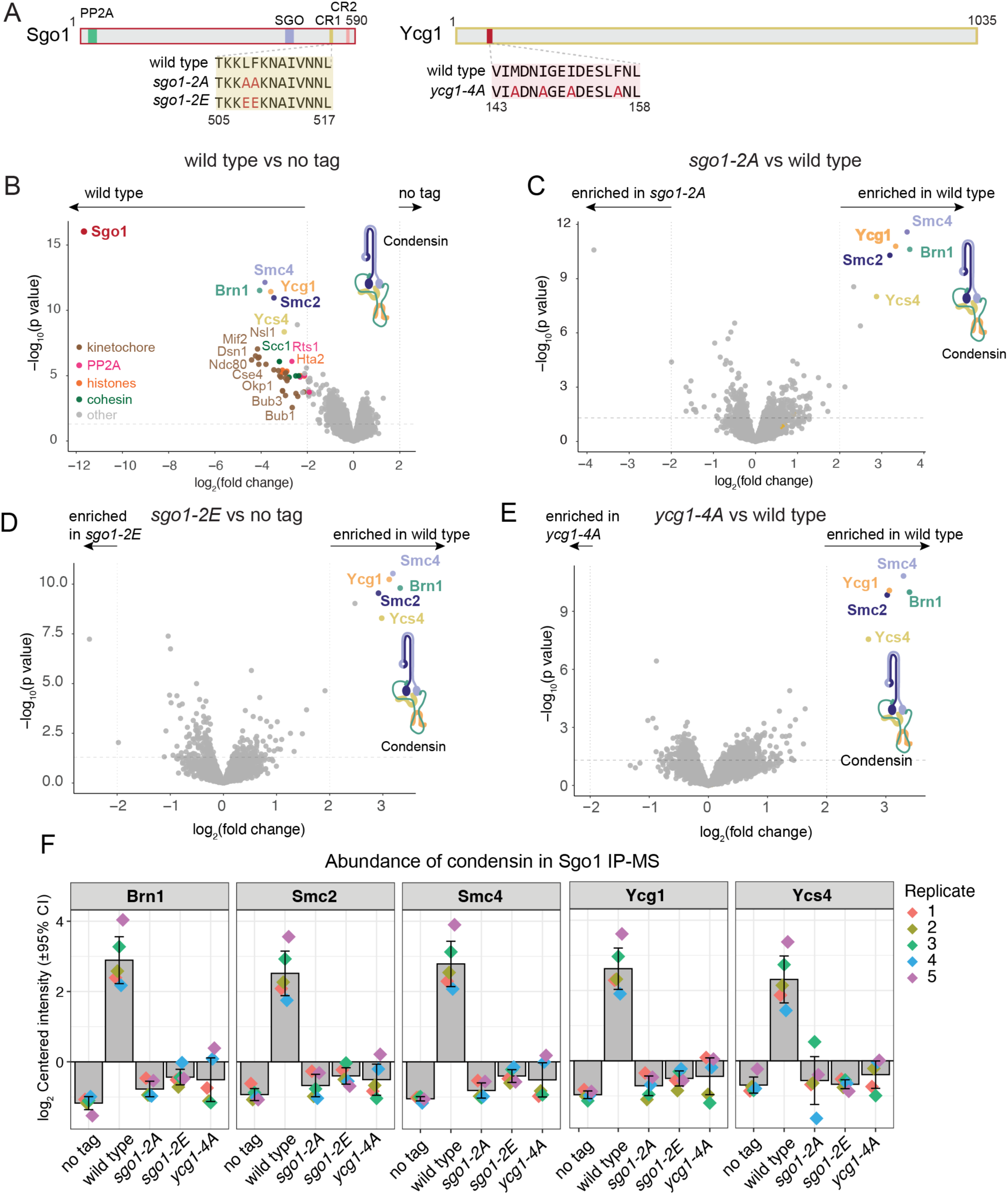
Sgo1-CR1 binding to Ycg1 occurs through the conserved pocket *in vivo*. (A) Scheme showing the endogenous *SGO1* and *YCG1* point mutants generated. (B-F) Sgo1-6His-3FLAG was immunoprecipitated from mitotically-arrested cells (by benomyl treatment) and the immunoprecipitates were analysed by mass spectrometry. Volcano plots showing the relative enrichment of proteins immunoprecipitated with wild type vs no tag (B), *sgo1-2A* vs wild type (C), *sgo1-2E* vs wild type (D) and *ycg1-4A* vs wild type (E). (F) Abundance of condensin subunits in Sgo1-FLAG IP-MS. Data represents values from 5 biological replicates. See also Figure S5.

### Multiple condensin ligands may use CR1-like motifs to bind Ycg1/CAP-G

Interestingly, the residues within the Sgo1 CR1 motif involved in binding to Ycg1 are reminiscent of the sequence in the tail of Kif4A previously found to be required for association with condensin^43,44^ (Figure S6A). This suggested that multiple ligands may associate with the conserved Ycg1/CAP-G pocket in a similar way to Sgo1. Consistently, although *ycg1-5A* (M145A, I148A, I151A, F156A, F185A) mutants are viable, indicating that the essential condensin function is intact, they form smaller colonies than either wild type or sgo1*-2E* mutants (Figure S6B), suggesting that binding of other ligands to the Ycg1 hydrophobic pocket influences growth. One candidate Ycg1 ligand is Lrs4, which is required for condensin positioning at the rDNA and mating locus and which we found to harbor a conserved CR1-like motif (Figure S6C).

To identify further potential condensin ligands that could associate with the conserved Ycg1/CAP-G patch we scanned proteins of interest for a consensus CR1-like motifs derived from comparing Sgo1 and Lrs4 ([KR]-[KR]-L-[FYVIT]-[KR]-x(1,3)-[IV]). This revealed the presence of a CR1-like motif in several chromosome-associated proteins, providing candidate condensin ligands for future studies (Figure S6D). Together, these observations raise the interesting possibility that multiple ligands use CR1-like motifs to dock onto the conserved Ycg1/CAP-G binding pocket.

### Condensin localization at pericentromeres requires direct interaction with Sgo1

During metaphase, prior to the establishment of tension-generating kinetochore-microtubule interactions, condensin associates with pericentromeres, but is not detected to significant levels along chromosome arms^24,30^. Like condensin, Sgo1 is also specifically localized to pericentromeres that are not under tension^20,29^, and indeed the pericentromeric localization of condensin depends on Sgo1^24,33^. We sought to understand whether direct Sgo1-Ycg1 binding is responsible for condensin recruitment to pericentromeres. First, we confirmed that *sgo1-2A, sgo1-2E* and *ycg1-4A* mutations which disrupt the Sgo1-Ycg1 interface do not greatly perturb the localization of Sgo1 itself. We performed calibrated ChIP-Seq to examine the chromosomal localization of Sgo1-6His-3FLAG, or its mutant variants, in metaphase-arrested cells (using the microtubule-depolymerizing drug nocodazole to prevent the establishment of kinetochore-microtubule attachments). With the exception of Sgo1-2E, which showed a general reduction in its association with chromatin, we observed a comparable pattern of Sgo1 localization in wild type and mutant cells (Figure 6A-C). In all cases, Sgo1 displayed characteristic twin peaks at centromeres, enrichment throughout the pericentromere and peaks at the flanking pericentromeric borders (Figure 6A-C). Therefore, consistent with the specific effects of the mutations on the Sgo1-Ycg1 interaction *in vivo* (Figure 5), disruption of the Sgo1-Ycg1 interface has little effect on the pericentromeric localization of Sgo1. Next we analysed the localization of Brn1-6HA by calibrated ChIP-Seq. As expected^24^, in wild type cells, there was a prominent Brn1 peak at all centromeres and, although the peak height varied between chromosomes, Brn1 was also detected at the pericentromeric borders that flank centromeres^20^ (Figure 6D and E). In contrast, in *sgo1-*2A, *sgo1-2E* and *ycg1-4A* mutants, condensin was lost throughout pericentromeres, except for at core centromeres, where a small peak remained, similar to *sgo11′* cells (Figure 6D-H). The mutations that perturb the Sgo1-Ycg1 interface show similar Brn1 localization to cells completely lacking Sgo1, indicating that disruption of this interface is sufficient to prevent Sgo1-dependent recruitment of condensin to pericentromeres. Interestingly, however, a modest Brn1 peak persisted at centromeres in all mutants, even *sgo1Δ* (Figure 6F). On closer inspection, we noticed only a single Brn1 peak in the mutants, in contrast to the characteristic bimodal peak of wild-type cells (Figure 6G). This could indicate the existence of a condensin recruiter in the kinetochore, that functions through a binding surface distinct from the Ycg1 hydrophobic pocket we found to bind Sgo1. Potentially the key function of Sgo1 is to relay kinetochore-recruited condensin to the pericentromere and borders, which critically depend on Sgo1 and the Sgo1-Ycg1 interface for condensin recruitment (Figure 6H). We conclude that condensin recruitment to pericentromeres is mediated through a direct interaction with Sgo1.

**Figure 6.**
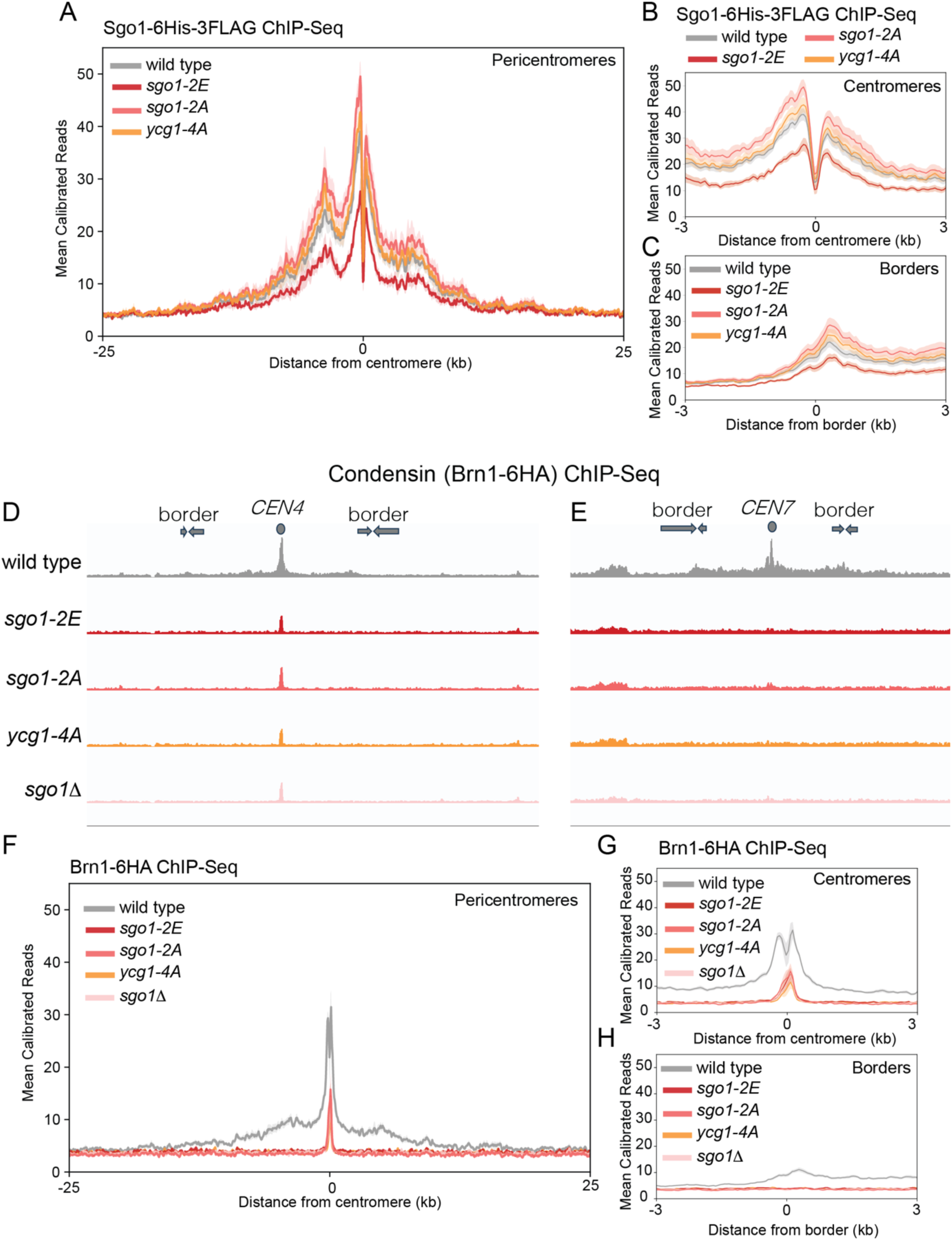
Sgo1 recruits condensin to pericentromeres through their direct interaction. (A-C) Calibrated Sgo1-6His-3FLAG ChIP-seq using cells arrested in metaphase by treatment with nocodazole. (A) Pileup of pericentromeric region of all 16 chromosomes. (B, C) Zoomed in pileups of a 6kb region surrounding 16 centromeres (B) or 32 pericentromeric borders (C). (D-H) Calibrated condensin (Brn1-6HA) ChIP-seq of cells arrested in metaphase by treatment with nocodazole. Condensin enrichment along representative sections of chromosome IV (D) and chromosome VII (E) including pericentromeres are shown. Pile-ups shows the mean ChIP-seq reads (solid line) with standard error (shading) at 16 pericentromeres (F) or zoomed in centromeres (G) and 32 borders (H).

### Pericentromeric condensin facilitates sister kinetochore biorientation in mitosis

Sister kinetochore biorientation in mitosis relies on Sgo1^24,26,54^. However, Sgo1 recruits several effectors to pericentromeres including Aurora B, PP2A and condensin^24,25^, and the extent to which Sgo1 promotes biorientation through condensin remains unclear. The availability of Sgo1 mutants that specifically abolish the interaction with condensin but not other binding partners (Figure 5) allowed us to ask if the Sgo1-Ycg1 interaction is important for sister kinetochore biorientation in mitosis. To test this, cells were released from a G1 arrest into a metaphase arrest (by withholding expression of the anaphase-promoting protein Cdc20) in the presence of nocodazole to depolymerize microtubules. Nocodazole was then washed out to allow spindle formation, while the metaphase arrest was maintained. These cells also carried a GFP label integrated close to the centromere of chromosome IV (*CEN4-* GFP) and a spindle pole body label (Spc42-tdTomato). As microtubules reform, sister kinetochore biorientation can be established, resulting in sufficient tension to split sister chromatid *CEN4-*GFP labels (Figure 7A). We scored the percentage of cells with two GFP foci through time after nocodazole washout as a measure of the efficiency and magnitude of sister kinetochore biorientation in the population, with the percentage of cells with two Spc42-tdTomato foci serving as a control for the numbers of metaphase cells in all conditions (Figure 7B). This revealed that the biorientation of sister kinetochores was both delayed and incomplete in *sgo1-2E* and *ycg1-4A* cells, albeit not to the extent of *sgo11′* cells (Figure 7C). Therefore, Sgo1 promotes sister kinetochore biorientation, in part through recruitment of condensin to pericentromeres via direct interaction with Ycg1.

**Figure 7.**
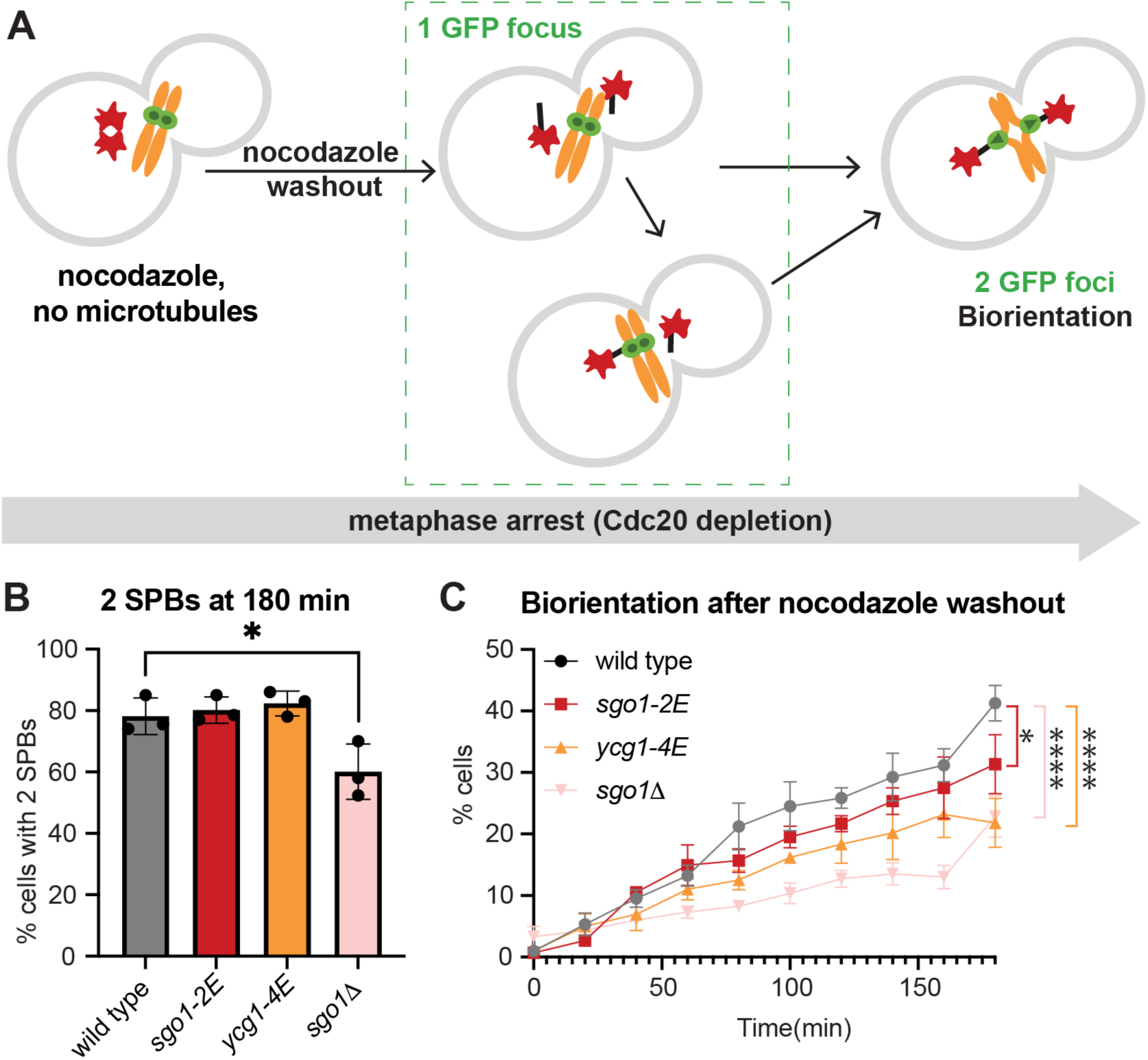
Sgo1 biases sister kinetochores to biorient by condensin recruitment. (A) Schematic diagram showing the biorientation assay. Wild type, *sgo1-2E*, *ycg1-4A* and *sgo1Δ* cells carrying *CEN4-GFP* (green) and SPB (*SPC42-tdTomato*, red) markers were released from a G1 arrest into nocodazole, and arrested in metaphase by Cdc20 depletion. After 3 hours, nocodazole was washed out (*t =* 0) and *CEN4-*GFP separation was scored every 20 mins for 3 hours. (B) Bar chart showing the percentage of cells with 2 SPB dots at the last timepoint (180 mins). Error bars represent deviation. *p<0.0332, one-way ordinary ANOVA with Dunnetts correction, only comparisons significantly different from wild type are indicated. (C) The percentage of separated *CEN4-GFP* foci across the timecourse are shown. Error bars represent standard error. p values refer to analysis of the last time point, *p<0.0332, ****p<0.0001, two-way ANOVA, with Tukey correction, only comparisons significantly different from wild type are indicated.

## Discussion

We have presented a molecular explanation for condensin enrichment at pericentromeres in budding yeast. We demonstrate that the HAWK subunit Ycg1 uses a conserved binding pocket to dock onto a conserved motif, CR1, on the pericentromeric adaptor protein, shugoshin, and that this is important for condensin localization at pericentromeres. We show that pericentromeric condensin promotes the efficient biorientation of sister kinetochores at mitosis. Our findings have implications for how condensin engages with ligands more generally. The binding pocket we identified on Ycg1 is broadly conserved and we discovered CR1-like motifs in the known condensin ligands mammalian KIF4A and yeast Lrs4, together with several potential ligands that we identified by homology search of the yeast proteome. While future work is required to test whether these candidates are true condensin ligands and to determine their functions, it is likely that additional chromosomal receptors use the same mode of condensin binding to define the architecture of chromatin sub-domains.

### Specific targeting of SMC proteins shapes the architecture of pericentromeres

The structuring of pericentromeres by cohesin and condensin allows them to perform specialized functions in chromosome segregation^15^. Our explanation of the specific targeting of condensin by Sgo1 adds to our understanding of how pericentromere architecture is defined. Initially, cohesin is recruited to kinetochores by Ctf19, resulting in the formation of a loop on either side of the centromere^20,21^. Sgo1, in turn, requires cohesin, as well as Bub1-dependent phosphorylation of H2A-S121 (Thr120 in humans) for its pericentromeric localization^24,29,49,55,56^. Finally, condensin docks onto Sgo1 to promote proper sister kinetochore biorientation. Interestingly, upon the establishment of biorientation, loop-extruding cohesin, Sgo1 and condensin are all released from pericentromeres, while cohesive cohesin is retained at pericentromere borders to resist pulling forces^20,29,30^.

How might pericentromeric condensin promote sister kinetochore biorientation? Chromosome capture experiments in budding yeast and vertebrates have observed structural changes at centromeres in cohesin-deficient cells^7,10^. Consistently, several studies have suggested that condensin provides a rigidity that allows pericentromeres to resist pulling forces and establish tension, which in turn is required to prevent kinetochore-microtubule destabilization by the error correction machinery^9,11,14,57^. An alternative, but not mutually exclusive possibility, is that condensin provides a geometry to pericentromeres that positions sister kinetochores in a back-to-back orientation, providing an intrinsic bias for sister kinetochores to attach to microtubules from opposite poles. In budding yeast, where a single microtubule contacts each kinetochore, such a bias has been observed to exist and requires both Sgo1 and condensin, in support of this idea^24,54,58^. Understanding the interplay between cohesin and condensin in defining pericentromere structure before and after tension establishment and how facilitates chromosome segregation is an important question for the future.

### Commonalities in regulation of SMC proteins by ligand-binding HAWKs

Our discovery of a conserved ligand-binding pocket on the condensin HAWK Ycg1, is reminiscent of the Conserved Essential Surface (CES) on cohesin. The CES is formed of a composite surface between the HAWK, SA2, and the kleisin RAD21 and is a binding site for multiple regulators^59,60^. These include the loop anchor protein CTCF, the cohesin release factor, WAPL1 and SGO1, which counteracts WAPL-dependent cohesin release^59,60^. Interestingly, the CES occupies a similar position on the SA2 to the conserved pocket on Ycg1 we identified here, suggesting that general principles may regulate SMC protein complexes.

### Limitations of the study

To predict the Ycg1-Sgo1 binding site, AlphaFold was used, which may not accurately represent the structure. However, the consistency of the model with our CLMS data and the demonstration that mutations in this interface specifically disrupt this interaction *in vivo* temper this concern. We focused on the interaction between a region in the C-terminal domain of Sgo1 and the N-terminal domain of Ycg1 and showed that mutations in this interface were sufficient to abrogate Sgo1 binding to condensin. However other parts of Sgo1, particularly in the N terminal region, may also contribute to binding. A previous study using peptide arrays found that condensin binding required a Serine-Rich Motif in Sgo1 residues 137-163, though this was not sufficient^33^. Our CLMS provided no evidence for interactions between condensin and Sgo1(137-163) and deletion of this region did not prevent Brn1 recruitment in the *in vivo* tethering assay. The reasons for this discrepancy remain unclear but could indicate an indirect role for residues Sgo1(137-163).

## Acknowledgements

We gratefully acknowledge the Wellcome Discovery Research Platform for Hidden Cell Biology Proteomics Core and Bioinformatics Core for mass spectrometry and bioinformatics support, respectively. We thank the Edinburgh Protein Purification Facility and Martin Wear for access to equipment. We thank Christian Häring, Damien D’Amours for plasmids and yeast strains. We are grateful to Owen Davies, Marcus Hassler, Christian Häring, Daniel Panne, Martin Singleton and Marcus Wilson for helpful discussions, and to Owen Davies, Alexander Julner Dunn, Lori Koch, Lucia Massari and Hollie Rowlands for comments on the manuscript. This work was funded through a Wellcome Investigator award to ALM [220780], two Wellcome Multi-User Equipment Grants [108504 and 2183052], core funding for the Wellcome Centre for Cell Biology [203149] and a Wellcome Discovery Research Platform Award [226791].

## Author Contributions

Conceptualization – MW and AM; Data curation –MW, JZ and CS; Funding Acquisition – AM; Formal Analysis – MW, JZ and CS; Investigation – MW; Supervision – AM and JR; Visualization – MW and AM; Writing – original draft – MW and AM; Writing – review and editing – MW and AM with input from all authors.

## Supplementary Information

**Table S1. Crosslink length within condensin subunits**

This table contains 3 sheets:

- The distance of crosslinks within condensin subunits BS3 CLMS data of full length Sgo1 with the pentameric condensin (Figure 1D) mapped onto cryo-EM of tetrameric condensin (lacking Ycg1; PBD:6YVU^46^; Figure S1B).
- The distance of crosslinks between Ycg1 and Brn1 BS3 CLMS of Sgo1(350-590) with Ycg1(6-932, Δ499-555)-Brn1(384-529) (Figure 2B) mapped onto a crystal structure (PBD:5OQQ^47^; Figure S2B)
- The distance of crosslinks between Ycg1 and Brn1 EDC CLMS of Sgo1(350-590) with Ycg1(6-932, Δ499-555)-Brn1(384-529) (data not shown) maps onto a reported crystal structure (PBD:5OQQ^47^; Figure S2B)

**Table S2. Table showing the distance of crosslinks between Sgo1 and Ycg1 for all five AlphaFold2 models**

**Figure S1.**
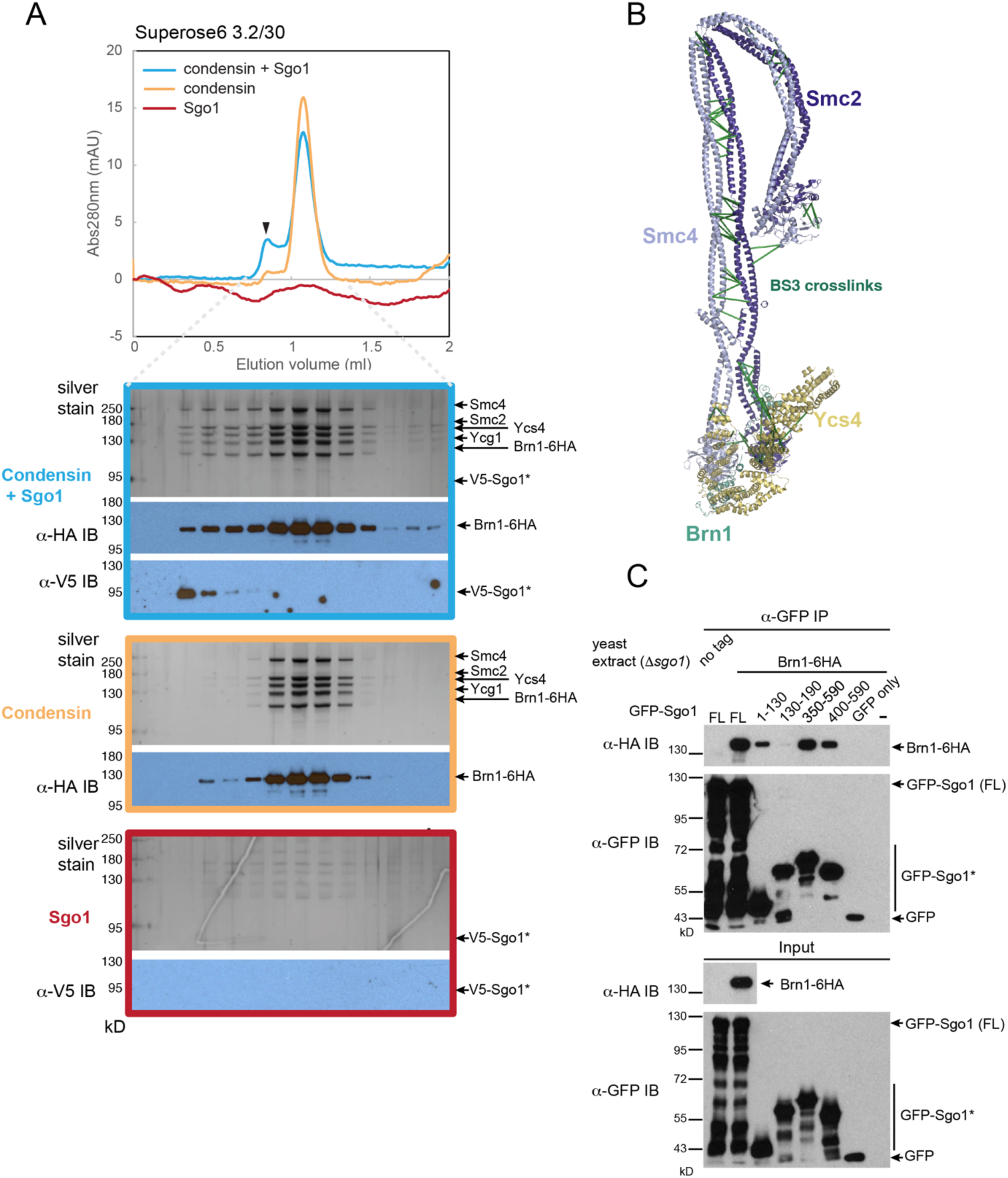
Analysis of Sgo1-condensin complexes. Related to Figure 1. (A) Size Exclusion Chromatography (SEC) profiles and corresponding silver-stained SDS-PAGE gels and immunoblot using the indicated antibodies for the analysis of full length Sgo1 (red) and condensin (yellow) complex formation (blue). Note that Sgo1 alone could not readily be detected in this assay as it associates non-specifically with the column in the absence of condensin. (B) Crosslinking mass spectrometry of full length Sgo1 with condensin data mapped onto reported condensin cryo-EM structure (PDB 6YVU^46^). Self-links are hidden and only crosslinks with score>10.5 are shown. (C) Purified GFP tagged Sgo1 variants co-immunoprecipitated with condensin (Brn1-6HA) from *sgo1Δ* yeast extract. Elutes were analysed by immunoblot with the indicated antibodies.

**Figure S2.**
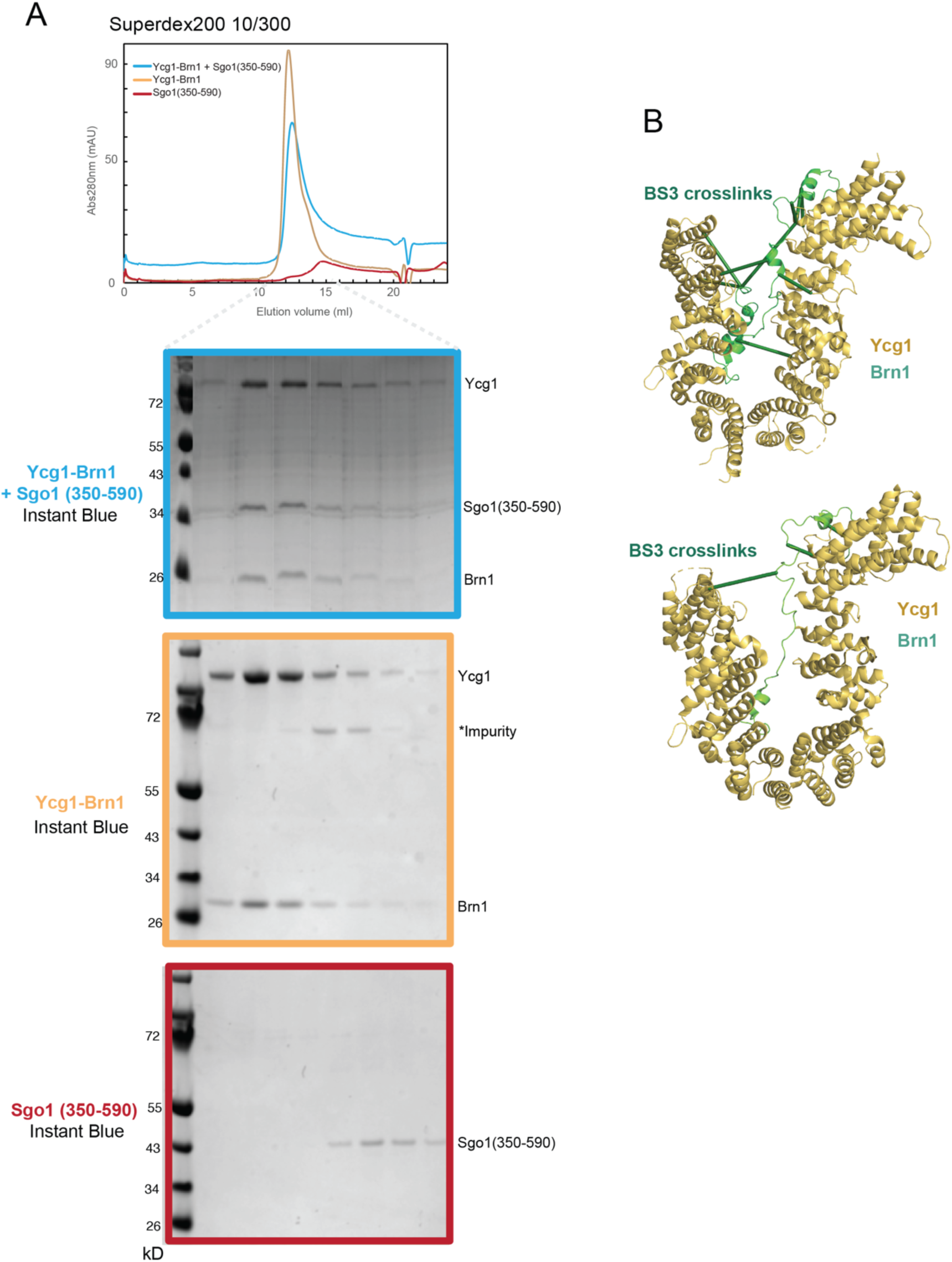
Analysis of C-Sgo1-Ycg1 complexes. Related to Figure 2. (A) SEC profiles and corresponding SDS-PAGE for the analysis of interactions between Sgo1 (350-590) and Ycg1 (6-932, Δ499-555)-Brn1 (384-529). (B) Crosslinking mass spectrometry of Sgo1 (350-590) and Ycg1 (6-932, Δ499-555)-Brn1 (384-529) mapped onto Ycg1-short Brn1 crystal structure which is a dimer (PDB 5OQQ^47^). Only crosslinks with score>10.5 were shown. Related to Figure 2

**Figure S3.**
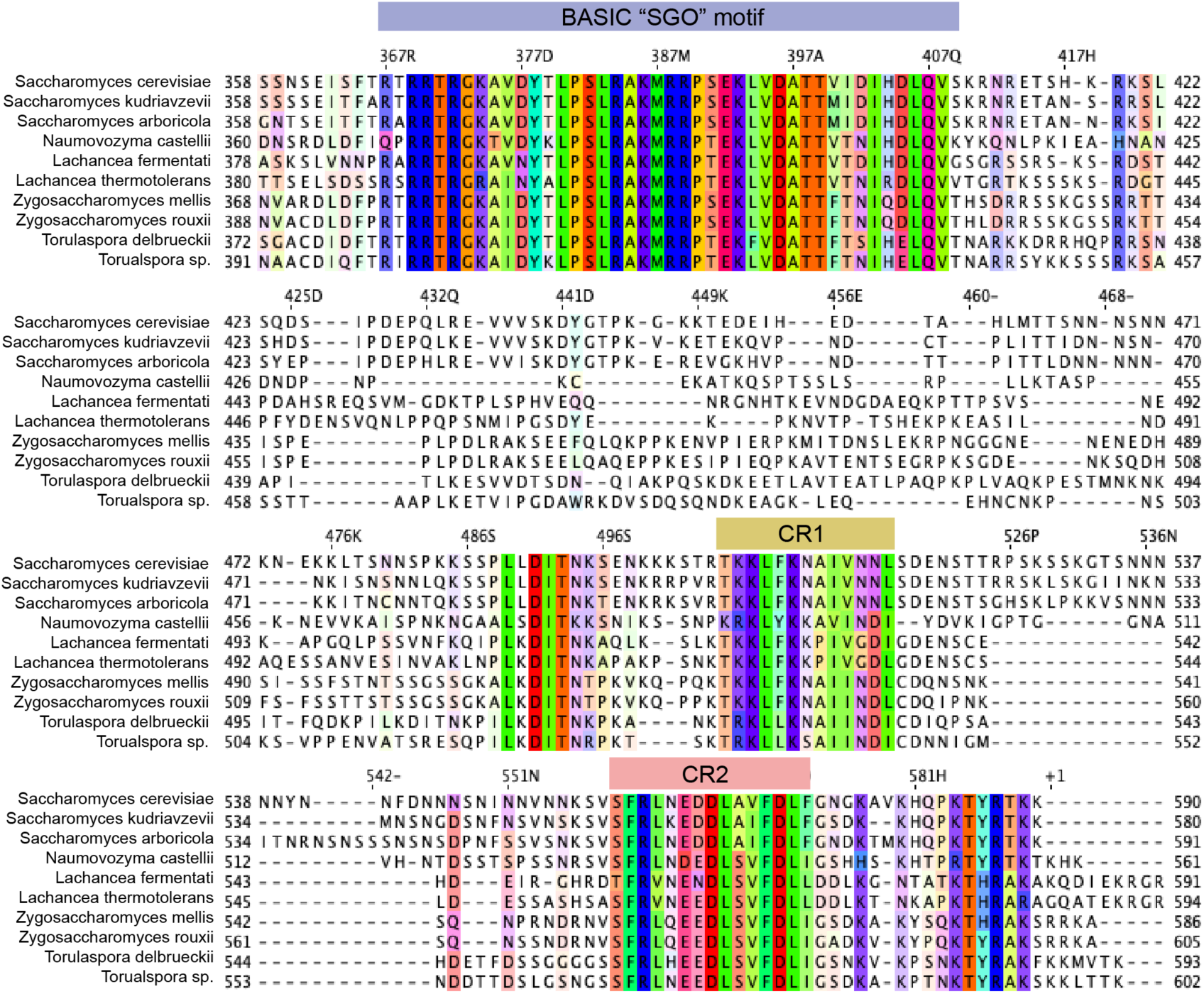
Conserved Regions in the C-terminal region of Sgo1. Related to Figure 3. Sequence alignment of C-terminus of Sgo1 among related yeast. The basic motif involved in histone binding is highlighted in blue. Two further conserved regions are highlighted in yellow and pink.

**Figure S4.**
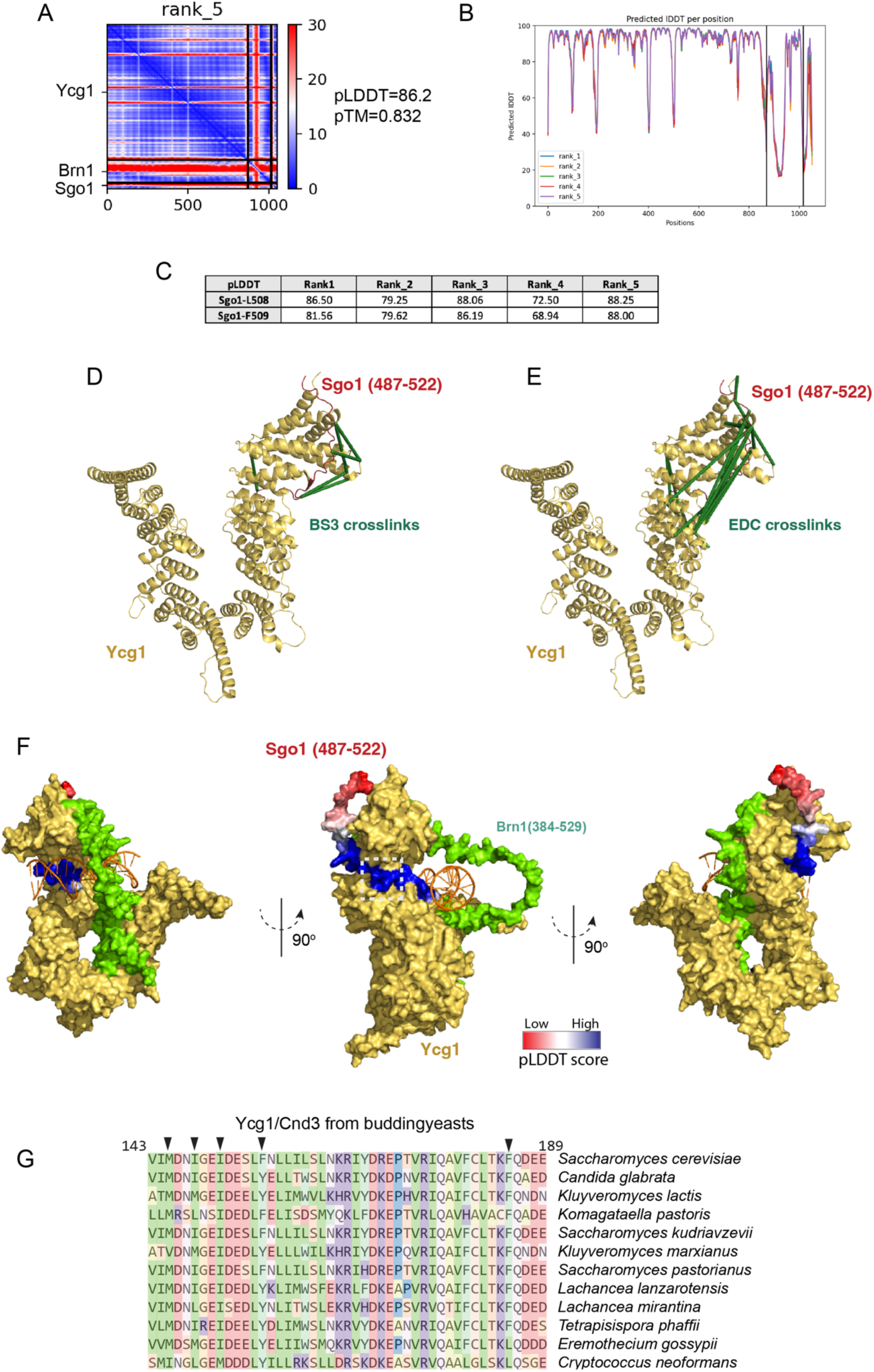
Agreement of the AlphaFold model with CLMS data. Related to Figure 4. (A) Predicted aligned error (PAE) matrices of AlphaFold2 model_Rank 5 obtained from the Sgo1(487-522) with Ycg1 (6-932, Δ499-555)-Brn1 (384-529) prediction (B) pLDDT plot of all 5 AlphaFold2 models. (C) Table showing the pLDDT confidence scores of Sgo1 residues L508, F509 for all 5 AlphaFold2 models. (D, E) Crosslinking mass spectrometry data of Sgo1 (350-590) and Ycg1 (6-932, Δ499-555)-Brn1 (384-529) with crosslinker BS3 (D) and EDC (E) map on AlphaFold2 model. (F) DNA was docked onto AlphaFold2 model by aligning with reported crystal structure (PDB 5OQP^47^). Sgo1 peptide (487-522) is coloured with pLDDT score. Sgo1-L508, F509 residues are highlighted in dashed box. (G) Conservation of the Ycg1 binding pocket in yeast.

**Figure S5.**
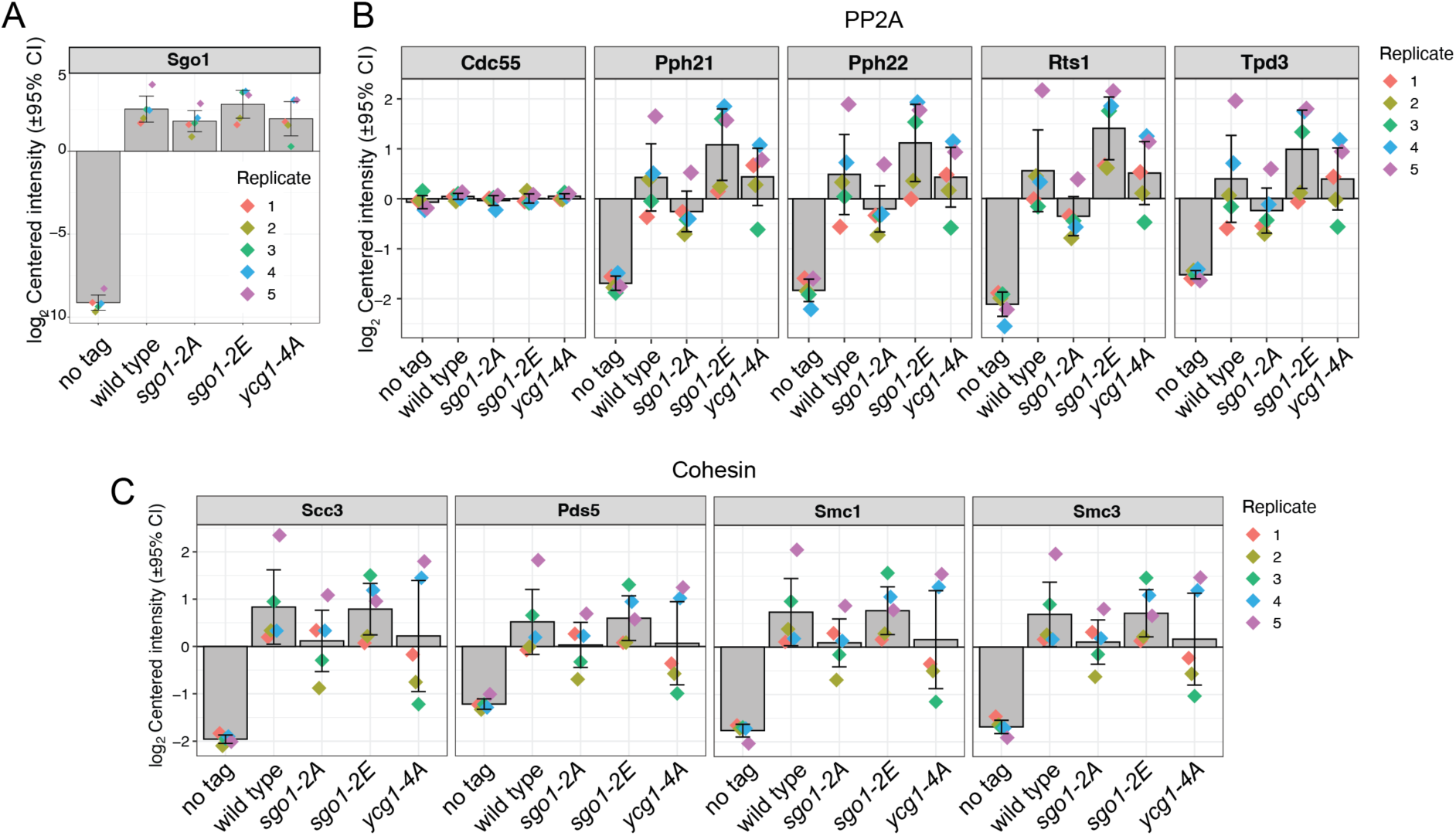
PP2A and cohesin association with Sgo1 is not affected by disruption of the CR1-Ycg1 interface. Related to Figure 5. Abundance of Sgo1 (A), PP2A (B), cohesin (C) in Sgo1-FLAG IP-MS.

**Figure S6.**
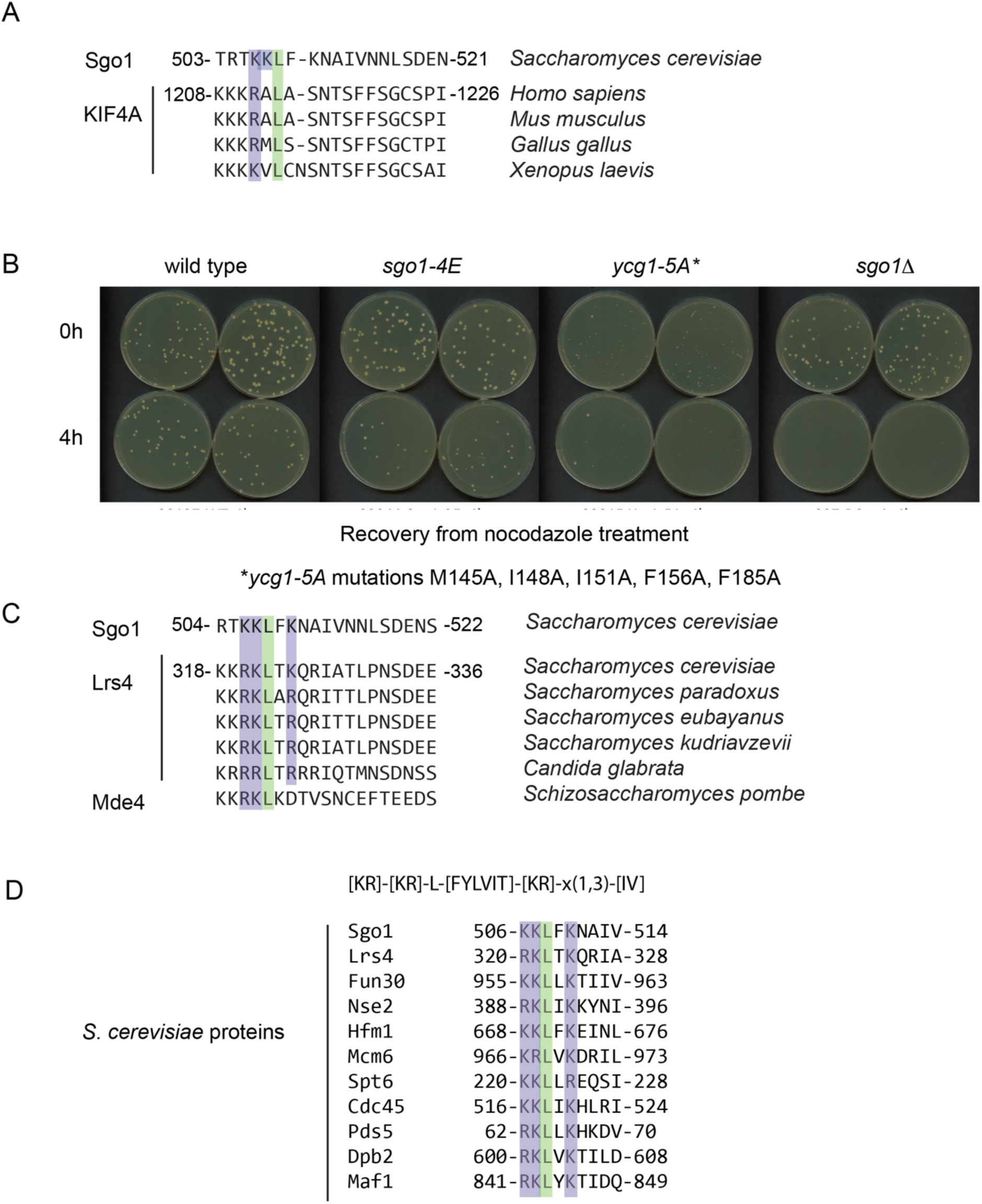
Identification of CR1-like motifs in other potential Ycg1/CAP-G ligands. Related to Figure 5. (A) A region of KIF4A that interacts with condensin I resembles the Sgo1 CR1. (B) Mutation of the Ycg1 binding pocket reduces colony size to a greater extent than mutation of the Sgo1-CR1. Cells were plated onto rich medium before (0h) or after addition of nocodazole (4h). (C) A CR1-like motif is found in the monopolin protein Lrs4/Mde4. (D) Potential candidate condensin ligands containing ([KR]-[KR]-L-[FYVIT]-[KR]-x(1,3)-[IV]) in *S. cerevisiae*.

## STAR methods

### RESOURCE AVAILABILITY

#### Lead contact

Further information and requests for resources and reagents should be directed to and will be fulfilled by the lead contact, Adele Marston

#### Materials availability

Yeast strains and plasmids used in this study can be obtained from the lead contact, without restriction.

#### Data and code availability

Mass spectrometry datasets reported in this study have been deposited at PRIDE and are publicly available as of the data of publication. DOIs are in the key resources table.

The paper does not report original code.

Any additional information required to reanalyze the data reported in this paper is available from the lead contact upon request.

### EXPERIMENTAL MODEL AND STUDY PARTICIPANT DETAILS

#### Plasmids

Plasmids used and generated in this study are listed in the Key Resources Table. *SGO1* fragments, deletion and point mutant variants were generated using PCR and cloned using Gibson assembly. *GFP-SGO1* used in in *E. coli* expression was codon optimized, and *V5-SGO1* used in yeast expression was described previously^45^. Constructs for generation of pentameric condensin complex and *YCG1 (6-932, Δ499-555) - BRN1 (384-529)* were kind gifts from Damien D’Amours^61^ and Christian Haering^47^.

#### Yeast strains

All yeast strains are W303 derivatives and are listed in the Key Resources Table. *SGO1-FLAG*, *BRN1-HA*, *CEN4-GFP* were described previously^24^. *SGO1* mutants were introduced using standard PCR-based methods. *YCG1-4A* was made by CRISPR-Cas9 in this study.

### METHOD DETAILS

#### IN VITRO METHODS

##### Protein expression and purification

All purified recombinant proteins are listed in the Key Resources Table. Expression and purification of V5-Sgo1 (FL) (used in co-immunoprecipitation and crosslinking mass spectrometry, Figure 1B-D, S1A) was described previously^45^. Briefly, protein was expressed in a protease-deficient yeast strain (AM8184) with 0.5 mM CuSO4 for 6h. After ball breaker grinding, cell powder was resuspended in lysis buffer (25 mM Tris-HCl, pH 7.5, 150 mM NaCl, 1 mM MgCl_2_, 10% glycerol, 0.1% NP-40, 0.05 mM EDTA, 0.05 mM EGTA, 1 mM DTT) plus protease inhibitors (1× CLAAPE [chymostatin, leupeptin, aprotinin, antipain, pepstatin, and E-64], 1 mM Pefabloc, 0.4 mM Na orthovanadate, 0.1 mM microcystin, 1 mM N-ethylmaleimide, 2 mM β-glycerophosphate, 1 mM Na pyrophosphate, 5 mM NaF and complete EDTA-free protease inhibitor (Roche)), and treated with 40 U/ml benzonase for 1.5 hours. The crude lysate was diluted with 25 mM Na phosphate, pH 7.5, 500 mM NaCl, 10% glycerol, 0.1% NP-40, 10 mM imidazole, 1 mM DTT and 0.25 mM PMSF, and after ultracentrifugation and filtering, loaded onto Ni^2+^conjugated HiTrap IMAC FF 1-ml column (GE Healthcare). After washing with 25 mM Na phosphate, pH 7.5, 500 mM NaCl, 10% glycerol, 0.1% NP-40, 25 mM imidazole, 1 mM DTT and 0.25 mM PMSF, protein was eluted with an increasing imidazole gradient (25–500 mM) over 40 column volumes and then loaded onto a gel filtration Superose 6 10/300 column in 50 mM Tris-HCl, pH 7.5, 500 mM NaCl, 10% glycerol, 0.1 mM EDTA, 0.1 mM EGTA, 1 mM DTT and 0.25 mM PMSF. V5-Sgo1 (350-590) (used in co-immunoprecipitation and crosslinking mass spectrometry, Figure 1E, 2A-B, S2A) was expressed in a protease-deficient yeast strain (AM8184) with 2% (w/v) galactose for 8h. The protein purification is the same as V5-Sgo1 (FL).

GFP-Sgo1 fragments (used in co-immunoprecipitation assays, Figure S1C) were expressed in *E. coli* BL21(DE3) with 0.1mM IPTG, at 18 °C overnight. Cells were resuspended in lysis buffer (20mM Na phosphate pH 8.0, 500mM NaCl, 1mM DTT, 10mM imidazole, 10% glycerol with complete EDTA-free protease inhibitor) after cell disruptor (25kPis) and ultracentrifuged, loaded onto Ni^2+^conjugated HiTrap IMAC FF 1-ml column (GE Healthcare) and elute with 25mM-500mM imidazole over 40 column volumes. The eluted protein was then loaded onto a gel filtration Superdex200 16/600 column in 50mM TrisHCl pH 8.0, 500mM NaCl, 1mM DTT, 0.25mM PMSF, 10% glycerol. For purification of GFP-Sgo1 C-terminus fragments, deletion and point mutant variants (used in co-immunoprecipitation assays, Figure 2C-G, 3B-C) purification, cells were resuspended in lysis buffer (25 mM HEPES, pH 7.5, 500 mM NaCl, 20mM imidazole, 1mM DTT, 10 % glycerol with complete EDTA-free protease inhibitor). After sonication and ultracentrifugation, lysates were incubated with Ni-NTA resin for 1 hour. The protein was then eluted with 200mM imidazole, and dialysed to no imidazole buffer. Pentameric condensin complex (purified from yeast, used in co-immunoprecipitation and crosslinking mass spectrometry, Figure 1B-E, S1A) and Ycg1 (6-932, Δ499-555) - Brn1 (384-529) (purified from *E. coli*, used in co-immunoprecipitation and crosslinking mass spectrometry, Figure 2A-B, S2A) were expressed and purified as described previously**^47^**.

#### Interaction studies using size exclusion chromatography

For full length Sgo1 and condensin holo-complex interaction studies (Figure S1A), Superose6 3.2/30 2.4ml column was used with buffer 20mM Tris pH 7.5, 200mM NaCl, 0.1mM EGTA, 1mM DTT and 0.05 ml fractions were collected. For Sgo1 (350-590) and Ycg1 (6-932, Δ499-555)-Brn1 (384-529) interaction studies (Figure S2A), Superdex200 10/300 24ml column was used, with buffer 10 mM TrisHCl pH 7.5, 250 mM NaCl, 5 % glycerol, 0.001% Tween20 and 0.3 ml fractions were collected.

#### Co-immunoprecipitation, western blotting, silver stain

##### Conjugating anti-V5, anti-GFP, anti-HA or anti-FLAG to dynabeads

Pre-washed (0.1M Na phosphate, pH 7.0) protein G dynabeads (Invitrogen) were incubated with anti-V5 (Biorad), anti-GFP (Roche), anti-HA (Biolegend) or anti-FLAG antibody (Sigma) with gentle agitation for 25 mins. After washing twice with 0.1M Na phosphate, pH 7.0, 0.01% Tween-20, then twice with 0.2 M triethanolamine, pH 8.2, proteins were cross linked with 20mM DMP (Dimethyl pimelimidate, Sigma) with rotation for 30 mins. After quenching with 50mM Tris-HCl, pH7.5, beads were washed three times with PBST (0.1% Tween-20).

##### Co-immunoprecipitation using two recombinant proteins (Figure 1BCE, 2ADF, 3B)

Pre-washed (25 mM HEPES pH 7.5, 500 mM NaCl, 10 % glycerol, 1 mM DTT, 0.1 % NP-40) antibody coupled dynabeads were incubated with two purified recombinant proteins with rotation for 2.5 hours. After washing 5 times with the same pre-wash buffer, proteins were eluted from the beads by boiling at 65 °C for 15 mins in 1x NuPAGE LDS sample buffer.

##### Co-immunoprecipitation using one recombinant protein and yeast extract (Figure S1C)

Cycling *sgo11′* yeast cells AM827 (no tag) and AM8834 (*BRN1-6HA*) were snap frozen as small ‘noodles’ and lysed by grinding. After resuspending with lysis buffer (25 mM HEPES, pH 7.5, 2 mM MgCl_2_ 15 % glycerol, 0.1% NP-40, 150 mM KCl, 0.1 mM EDTA, 0.5 mM EGTA) plus protease inhibitors (1× CLAAPE [chymostatin, leupeptin, aprotinin, antipain, pepstatin, and E-64], 1 mM Pefabloc, 0.4 mM Na orthovanadate, 0.1 mM microcystin, 1 mM N-ethylmaleimide, 2 mM β-glycerophosphate, 1 mM Na pyrophosphate, 5 mM NaF and complete EDTA-free protease inhibitor), the crude lysates were treated with 40 U/ml of benzonase (Novagen) or BaseMuncher (Abcam) for 1 hour and centrifuged at 3600 rpm for 20 mins. Pre-washed anti-GFP coupled beads were incubated with recombinant GFP-Sgo1 fragments with rotating for 1.5 hours, and then washed 3 times with lysis buffer plus 0.25 mM PMSF. These Sgo1-beads were then incubated with the yeast lysate supernatant (*BRN1-6HA*) for another 1.5 hours. Following washing 5 times with lysis buffer plus 0.25 mM PMSF, proteins were eluted from the beads by boiling at 65 °C for 15 mins in 1x NuPAGE LDS sample buffer.

##### Western blotting

Proteins were separated in SDS-PAGE gels and transferred to nitrocellulose membranes. All antibodies were diluted in 2% milk PBST, and are listed in the Key Resources Table. Signals were detected by Pico-ECL (Thermo Fisher Scientific) and autoradiograms.

##### Silver stain

Proteins were separated in NuPAGE 4-12% Bis-Tris gel (Invitrogen) and stained with SilverQuest Silver Staining Kit (Invitrogen).

#### Crosslinking Mass Spectrometry

For BS3 crosslinking of full length Sgo1 and condensin holo-complex (Figure 1D; Table S1), 1:1 (w:w) gel filtrated proteins were incubated 1 hour on ice, after buffer exchanging to 25 mM HEPES, pH7.5, 150 mM NaCl, 5% glycerol, 1mM DTT, protein complex was crosslinked with BS3 (Thermo Scientific, BS3:protein ratio=3:1, w:w) for 2 hours on ice. The crosslinking was quenched by 100 mM ammonia bicarbonate and was briefly resolved using a NuPAGE 3-8% Tris-Acetate gel (Invitrogen, EA0378BOX). Bands were visualised by quick InstantBlue staining (Abcam), excised, reduced with 8 mM TCEP for 20 min at room temperature, alkylated with 5 mM iodoacetamide for 20 min at room temperature, digested at 37 °C using GluC (Thermo, GluC:protein ratio=1:10, w:w in 10mM ammonia bicarbonate) overnight, and then with 13 ng/μl trypsin (Pierce) at 37 °C for another 8 hours.

For BS3 crosslinking of Sgo1 (350-590) and Ycg1 (6-932, Δ499-555)-Brn1 (384-529) (Figure 2B; TableS1), purified proteins (Sgo1: Ycg1-Brn1 molar ratio=1.5:1) were incubated 1 hour on ice in 25 mM HEPES pH7.5, 500 mM NaCl, 10% glycerol. After crosslinking with BS3 (Thermo Scientific, BS3:protein ratio=2:1, w:w) for 2 hours on ice, proteins were resolved using a NuPAGE 4-12% Bis-Tris gel (Invitrogen). Similar steps were performed as above, except proteins were digested only with 13 ng/μl trypsin (Pierce) at 37 °C overnight.

For EDC crosslinking of Sgo1 (350-590) and Ycg1 (6-932, Δ499-555)-Brn1 (384-529) (Figure S4E; Table S1), Sgo1: Ycg1-Brn1 molar ratio was 3:1. 10 ug of protein complex was crosslinked with 10ug EDC (Thermo Fisher Scientific) and 22ug NHS (Thermo Fisher Scientific) in 25 mM HEPES pH6.8, 500 mM NaCl, 10% glycerol for 1.5 hours at room temperature.

LC-MS/MS analysis for was performed on an Orbitrap Fusion Lumos (Thermo Fisher Scientific). Peptide separation was carried out on an EASY-Spray column (50 cm × 75 μm i.d., PepMap C18, 2 μm particles, 100 Å pore size, Thermo Fisher Scientific). Mobile phase A consisted of water and 0.1% formic acid. Mobile phase B consisted of 80% acetonitrile and 0.1% formic acid. Peptides were loaded onto the column with 2% B at 300 nL/min flow rate and eluted at 200 nL/min flow rate in two steps: linear increase from 2% B to 40% B in 109 minutes; then increase from 40% to 95% B in 11 minutes. The eluted peptides were directly sprayed into the mass spectrometer. Peptides were analysed using a high/high strategy: both MS spectra and MS2 spectra were acquired in the Orbitrap. MS spectra were recorded at a resolution of 120,000. The ions with a precursor charge state between 3+ and 8+ were isolated with a window size of 1.6 m/z and fragmented using high-energy collision dissociation (HCD) with collision energy 30. The fragmentation spectra were recorded in the Orbitrap with a resolution of 30,000. Dynamic exclusion was enabled with single repeat count and 60 s exclusion duration.

The mass spectrometric raw files were processed into peak lists using ProteoWizard (version 3.0)^62^, and crosslinked peptides were matched to spectra using xiSEARCH software (version 1.7.6.1 and 1.7.6.4)^63^ (https://github.com/Rappsilber-Laboratory/XiSearch) with preprocessing and in-search assignment of monoisotopic peaks^64^. Search parameters were MS accuracy, 3 ppm; MS/MS accuracy, 10 ppm; enzyme, trypsin; crosslinker, BS3 or EDC; max missed cleavages, 4; missing mono-isotopic peaks, 2; fixed modification, carbamidomethylation on cysteine; variable modifications, oxidation on methionine and phosphorylation on serine and threonine; fragments, b and y ions with loss of H2O, NH3 and CH3SOH. The mass spectrometry proteomics data will be^65^ deposited to the ProteomeXchange Consortium via the PRIDE partner repository upon publication.

##### AlphaFold Predictions and identification of CR1-like motifs

AlphaFold prediction was performed using Google Colab multimer^50,51,66^ with num_relax set to 5 using protein sequences Sgo1(487-522), Ycg1 (6-932 Δ499-555) and Brn1 (384-529). To identify potential Ycg1 interacting proteins we searched for motifs with the consensus [KR]-[KR]-L-[FYVIT]-[KR]-x(1,3)-[IV] in the yeast genome using the Genomenet tool (https://www.genome.jp/tools/motif/) and selected the chromosomal proteins of interest as shown Figure S6D.

#### IN VIVO METHODS

##### Yeast growth and culture

Alpha factor was used at 5 μg/ml and re-added to 2.5 μg/ml every 1.5 hours. Nocodazole was used at 15 μg/ml and re-added to 7.5 μg/ml every 1 hour. Benomyl was used at 30 μg/ml. Methionine was used at 8 mM and re-added to 4 mM every 45 mins.

##### Immunoprecipitation and mass spectrometry

###### Immunoprecipitation

1 litre of culture grown at OD_600_= 1.8 was treated with 30 μg/ml benomyl for 2 hours, and then harvested and drop frozen in liquid nitrogen. Cells were cryo-lysed with freezer mill (Spex 6875, 8 rounds of 2 mins at 10 cycles/second, with 2 mins rests). Immunoprecipitation was done using anti-FLAG coupled dynabeads as described in *Co-immunoprecipitation using yeast extract*. Proteins were eluted from beads in two rounds by incubating with 0.1% RapiGest (Waters) in 50 mM TrisHCl, pH 8.0 at 50 °C for 10 mins with 500 rpm shaking.

Protein samples from all biological replicates were processed at the same time and with using the same digestion protocol without any deviations. They were subjected for MS analysis under the same conditions. Protein and peptide lists were generated using the same software and the same parameters. Specifically, eluates from each IP were digested using the Filter Aided Sample Preparation (FASP) protocol as described)^67^ with minor modifications. In brief, IP eluate was reduced with 25 mM DTT at 80°C for 1 min, then denatured by addition of urea to 8M. Sample was applied to a Vivacon 30k MWCO spin filter (Sartorius,UK) and centrifuged at 12.5k g for 15-20 minutes. Protein retained on the column was then alkylated with 100 μL of 50 μM iodoacetamide (IAA) in buffer A (8 M urea, 100 mM Tris pH 8.0) in the dark at RT for 20 min. The column was then centrifuged as before, and washed with 100 μL buffer A, then with 2 x 100 μL volumes of 50 mM ammonium bicarbonate (ABC). 3 μg/μL trypsin (Pierce, UK) in 0.5 mM ABC were applied to the column, which was then incubated at 37° overnight. The eluates from the filter units were acidified using 20 µl of 10% Trifluoroacetic Acid (TFA) (Sigma Aldrich), and spun onto StageTips as described^68^. Peptides were eluted in 40 μL of 80% acetonitrile in 0.1% TFA and concentrated down to 1 μL by vacuum centrifugation (Concentrator 5301, Eppendorf, UK). The peptide sample was then prepared for LC-MS/MS analysis by diluting it to 5 μL by 0.1% TFA.

LC-MS analyses were performed on an Orbitrap Exploris™ 480 Mass Spectrometer (Thermo Fisher Scientific, UK) coupled on-line, to an Ultimate 3000 HPLC (Dionex, Thermo Fisher Scientific, UK). Peptides were separated on a 50 cm (2 µm particle size) EASY-Spray column (Thermo Scientific, UK), which was assembled on an EASY-Spray source (Thermo Scientific, UK) and operated constantly at 50°C. Mobile phase A consisted of 0.1% formic acid in LC-MS grade water and mobile phase B consisted of 80% acetonitrile and 0.1% formic acid. Peptides were loaded onto the column at a flow rate of 0.3 μL min^-1^ and eluted at a flow rate of 0.25 μL min^-1^ according to the following gradient: 2 to 40% mobile phase B in 150 min and then to 95% in 11 min. Mobile phase B was retained at 95% for 5 min and returned back to 2% a minute after until the end of the run (190 min).

Survey scans were recorded at 120,000 resolution (scan range 350-1650 m/z) with an ion target of 5.0e6, and injection time of 20ms. MS2 Data Independent Acquisition (DIA) was performed in the orbitrap at 30,000 resolution with a scan range of 350-1200 m/z, maximum injection time of 55ms and AGC target of 3.0E6 ions. We used HCD fragmentation^69^ with stepped collision energy of 25.5, 27 and 30. We used variable isolation windows throughout the scan range ranging from 10.5 to 50.5 m/z. Shorter isolation windows (10.5-18.5 m/z) were applied from 400-800 m/z and then gradually increased to 50.5 m/z until the end of the scan range. The default charge state was set to 3. Data for both survey and MS/MS scans were acquired in profile mode.

The DIA-NN software platform^70^ version 1.8.1. was used to process the raw files and search was conducted against the *Saccharomyces cerevisiae* complete/reference proteome (Uniprot, released in December, 2019). Precursor ion generation was based on the chosen protein database (automatically generated spectral library) with deep-learning based spectra, retention time and IMs prediction. Digestion mode was set to specific with trypsin allowing maximum of two missed cleavages. Carbamidomethylation of cysteine was set as fixed modification. Oxidation of methionine, and acetylation of the N-terminus and phosphorylation of threonine serine and tyrosine were set as variable modifications. The parameters for peptide length range, precursor charge range, precursor m/z range and fragment ion m/z range as well as other software parameters were used with their default values. The precursor FDR was set to 1%. Statistical analysis was performed by Perseus software, version 1.6.2.1^71^. Raw data will be available after publication on the PRIDE database http://www.ebi.ac.uk/pride_no.

##### Chromatin immunoprecipitation-qPCR (ChIP-qPCR)

ChIP and qPCR were performed as described previously using anti-HA 12CA5, anti-GFP or anti-FLAG antibody^45^. Primers used for qPCR analysis are given in the Key Resources Table, and the reactions were carried out on a LightCycler 480 machine (Roche) with 45 cycles for NEB Luna. Experiments were done with four repeats and error bars represent standard error.

##### Calibrated Chromatin Immunoprecipitation and sequencing (ChIP-Seq)

ChIP-Seq was done as described previously with minor modifications^20,72^. Shortly, 400 ml of nocodazole arrested yeast culture was grown and fixed as for ChIP-qPCR. To each condition, equal amount of *S. pombe* (using AMsp635 for Brn1-6HA ChIP-seq, and AMsp1863 for Sgo1-FLAG ChIP-seq) fixed cell pellets were added for calibration. Cells were then processed as for ChIP-qPCR except 3 rounds of 30 seconds on Fastprep (BioPulverizer FP120), followed by 2 rounds of sonication (20 cycles of 30 seconds on/30 seconds off, at HIGH setting, Bioruptor Plus Diagenode). For every 1ml IP, 15 μl of Protein G Dynabeads (Invitrogen) and 7.5 μl of 12CA5 anti-HA or 5 μl of anti-FLAG antibody were used.

ChIP-sequencing libraries were prepared using NEXTflex-6 DNA Barcodes (PerkinElmer), quantified by Qubit (Thermo Fisher Scientific), and then library quality was tested using the 2100 Bioanalyzer High Sensitivity DNA kit (Agilent, Santa Clara, CA) on Bioanalyzer (Agilent). 1 nM pooled library (INPUT:IP ratio = 15:85%) was sequenced in house on an Illumina MiniSeq instrument (Illumina, San Diego, CA) with MiniSeq High output reagent kit (150-cycles, paired -end) (Illumina).

Data analysis was performed as described previously^72^. Mean plots (pileup) at 16 centromeres and 32 pericentromeric borders sites were generated using the Deeptools computeMatrix and PlotProfle package. Reads were binned at 50 bp windows around midpoint of centromeres or borders with 3 or 25 kb flanks at either side. Borders were oriented so that their position relative to the centromere was the same. Centromere, pericentromere border peak coordinates are as described previously^20^.

##### Biorientation assay

Biorientation assay in Figure 7 was performed with minor modifications as previously described^24^. Briefly, cells carrying *pMET-CDC20*, *CEN4-GFP* and *SPC42-tdTomato* were arrested in G1 phase using alpha factor in medium lacking methionine for 3 hours, then released to rich medium (YPDA) containing methionine, nocodazole and benomyl. Methionine represses *pMET-CDC20* to achieve metaphase arrest, nocodazole and benomyl drugs depolymerize microtubules. After 3 hours treatment, t0 sample was taken and drugs were washed out and released into rich medium containing methionine to allow spindles reforming maintaining metaphase arrest. Samples were taken every 20 mins by fixing in 3.7% formaldehyde for 10 min and resuspended in 1xDAPI for microscopy. Typically 200 cells (at least 100 cells) were scored for each timepoint.

### QUANTIFICATION AND STATISTICAL ANALYSIS

Statistical analysis and graphs were generated using Graphpad Prism 10 software (San Diego). Micrographs and graphs were assembled using Adobe Illustrator. Sequence alignment was generated using JalView. IP-MS data was analyzed using R studio. AlphaFold model was generated using AlphaFold2. Details of replicates are given in the figure legends.

### KEY RESOURCES TABLE

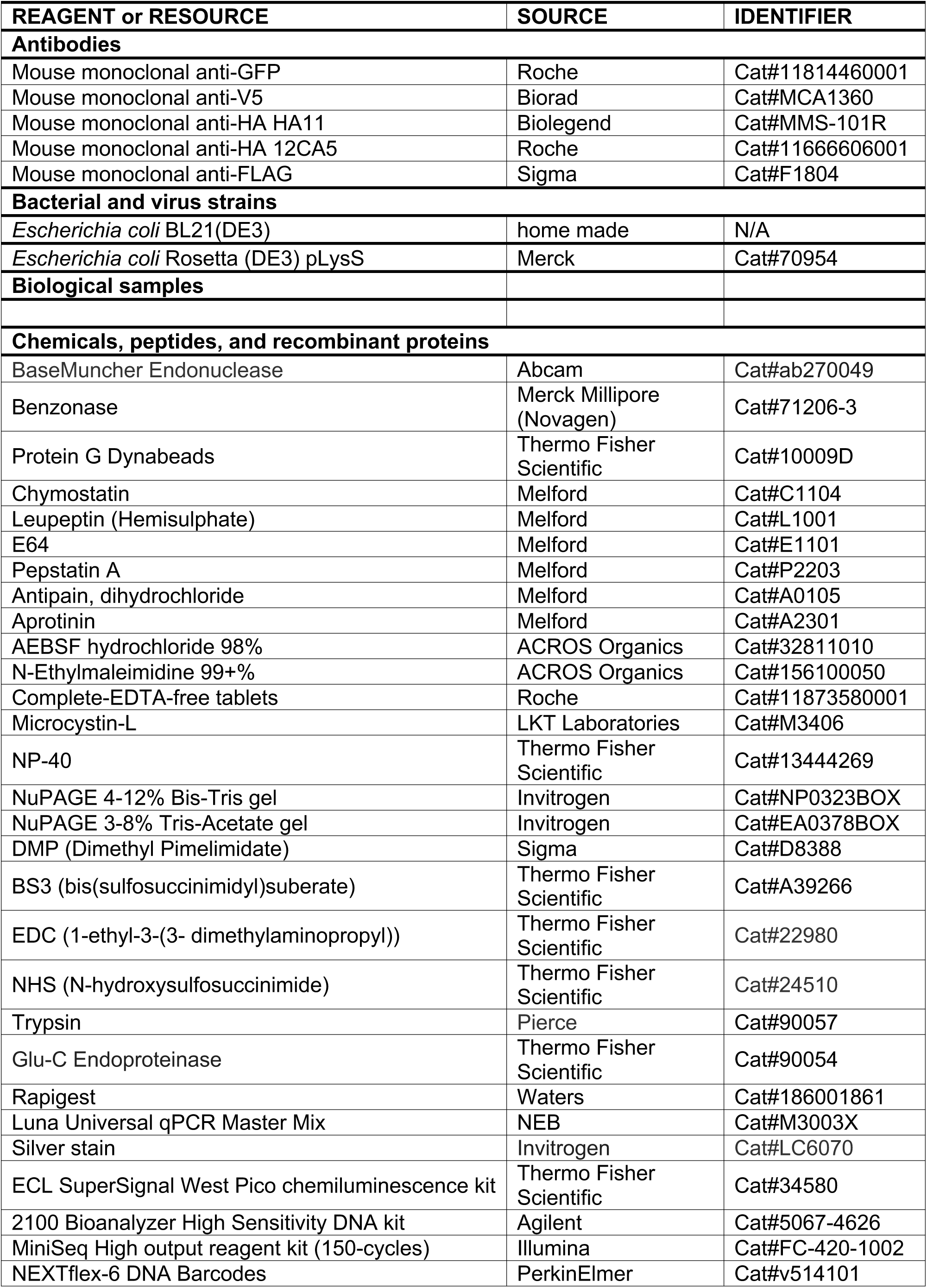
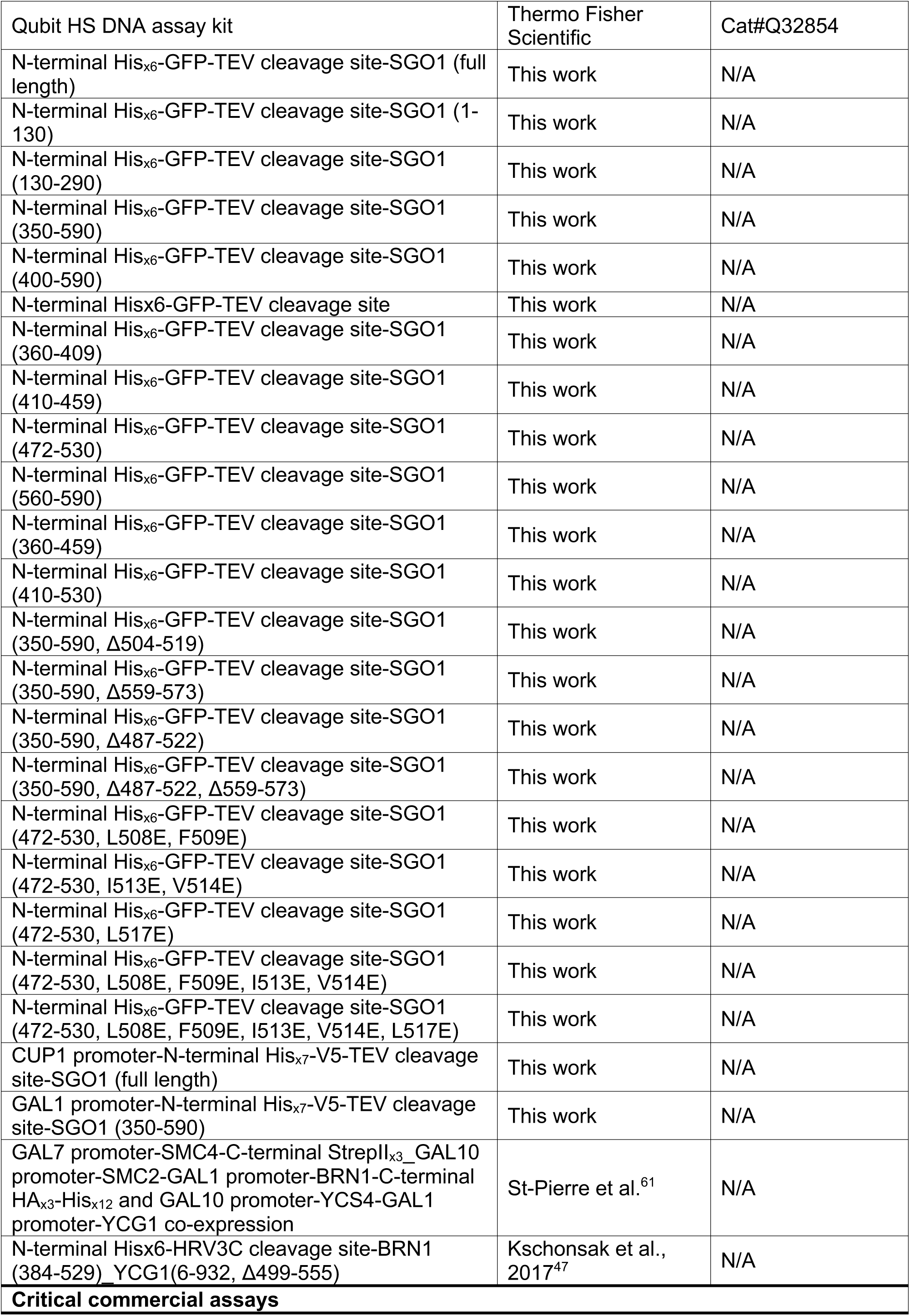
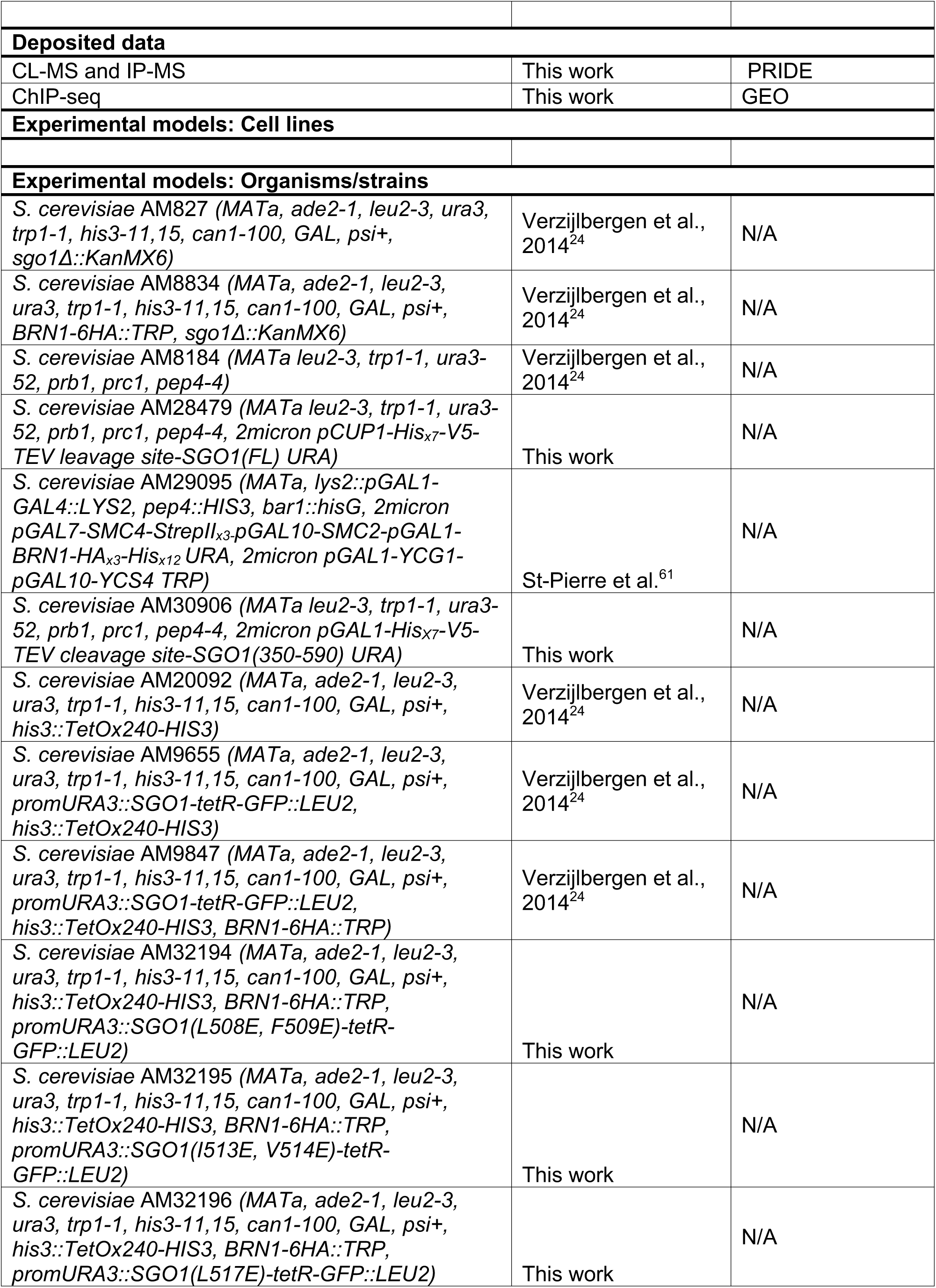
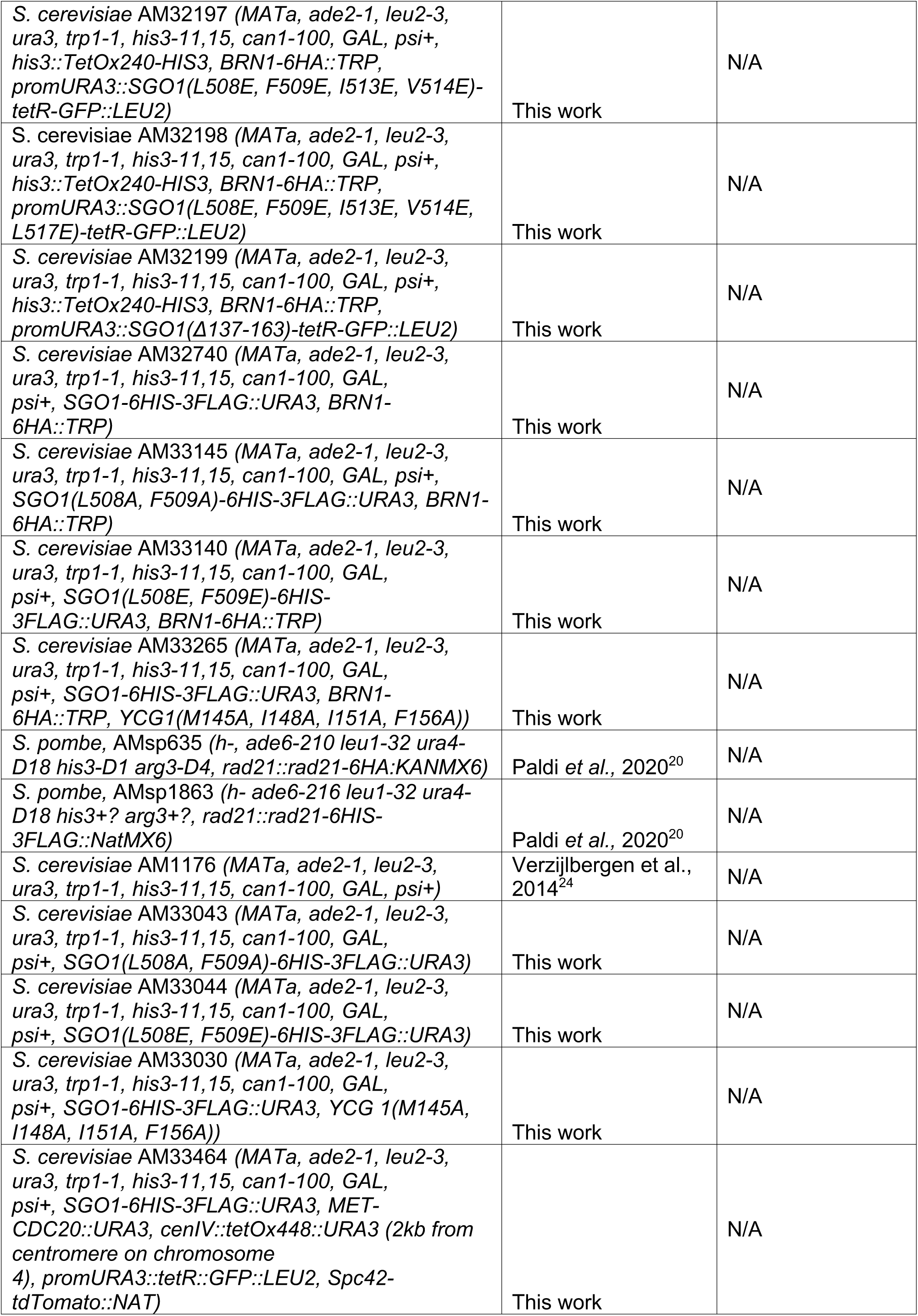
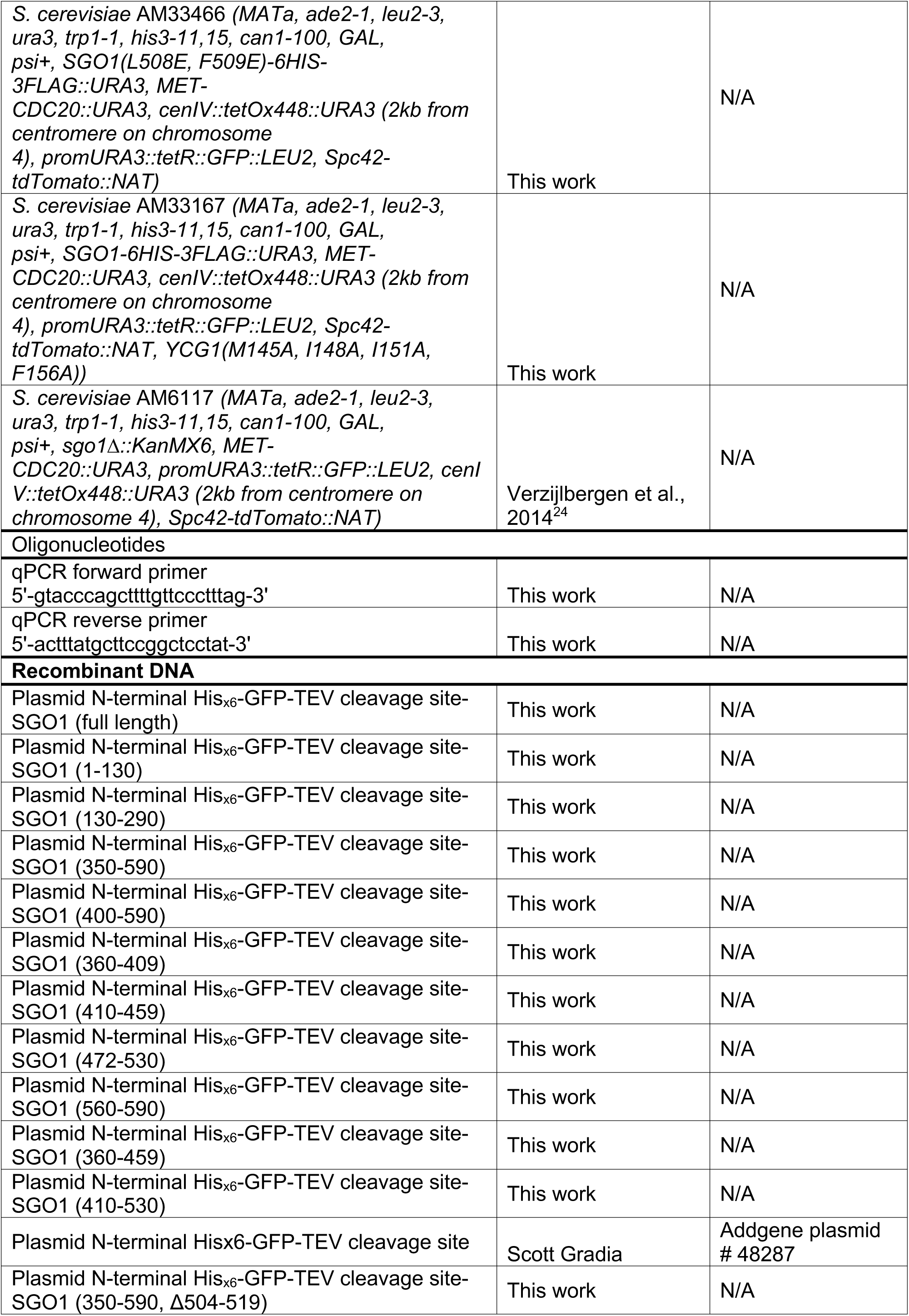
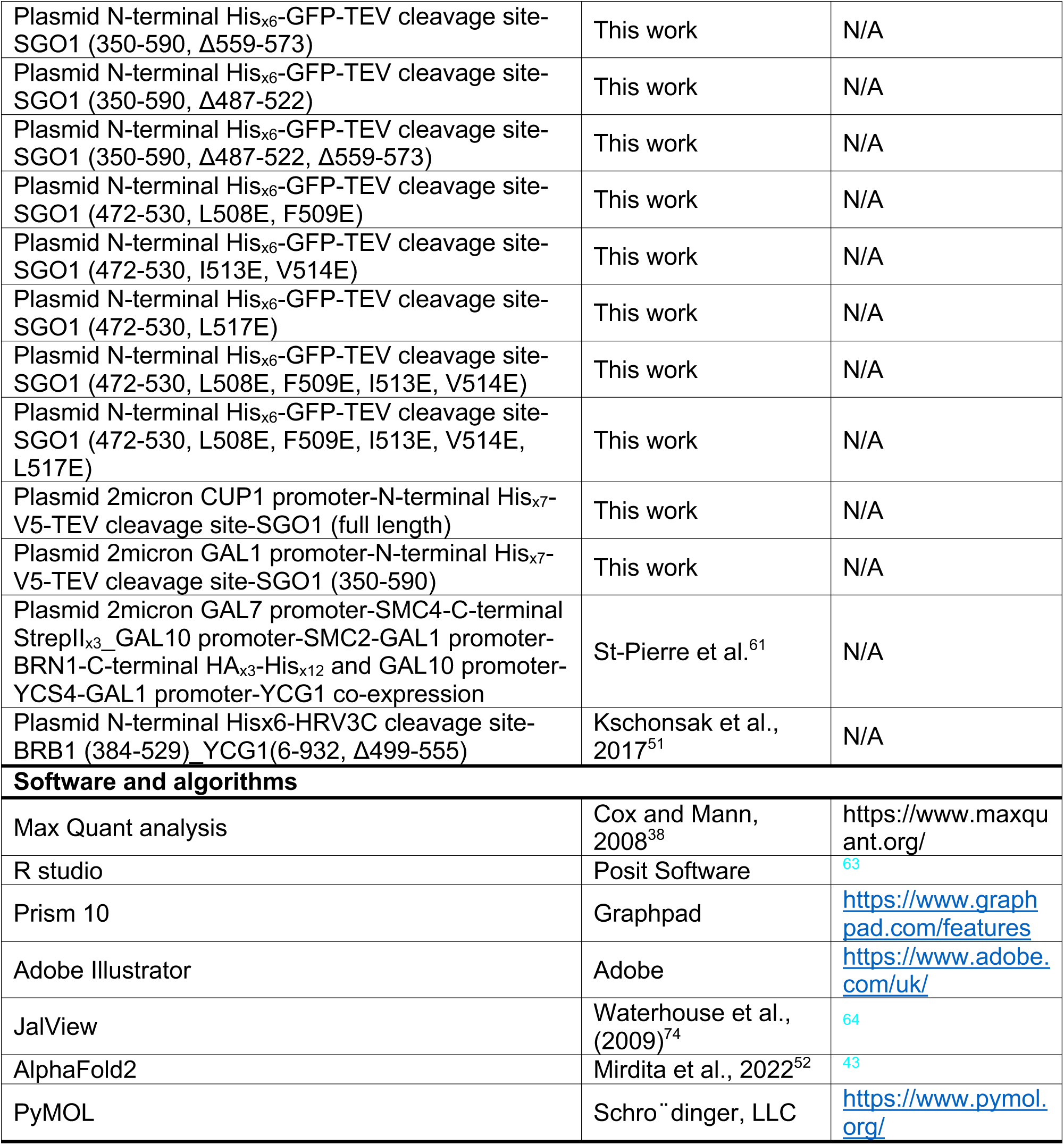

